# PPTC7 antagonizes mitophagy by promoting BNIP3 and NIX degradation via SCF^FBXL4^

**DOI:** 10.1101/2024.02.23.581814

**Authors:** Giang Thanh Nguyen-Dien, Brendan Townsend, Prajakta Kulkarni, Keri-Lyn Kozul, Soo Siang Ooi, Denaye Eldershaw, Saroja Weeratunga, Meihan Liu, Mathew JK Jones, S Sean Millard, Dominic CH Ng, Michele Pagano, David Komander, Michael Lazarou, Brett M Collins, Julia K Pagan

**Affiliations:** Faculty of Medicine, School of Biomedical Sciences, University of Queensland, Brisbane, QLD, Australia; Department of Biotechnology, School of Biotechnology, Viet Nam National University-International University, Ho Chi Minh City, Vietnam; The University of Queensland, Institute for Molecular Bioscience, Queensland, 4072, Australia; The University of Queensland Frazer Institute, Faculty of Medicine, The University of Queensland, Brisbane, QLD 4102, Australia; School of Chemistry & Molecular Biosciences, University of Queensland, Brisbane, QLD, 4072, Australia; Clem Jones Centre for Ageing Dementia Research, Queensland Brain Institute, The University of Queensland, Brisbane, QLD 4072, Australia; Department of Biochemistry and Molecular Pharmacology, New York University Grossman School of Medicine, New York, NY 10016, USA; Perlmutter Cancer Center, New York University Grossman School of Medicine, New York, NY 10016, USA; Department of Pathology and Lab Medicine, Meyer Cancer Center, Weill Cornell Medicine, New York, NY 10065, USA; Walter and Eliza Hall Institute of Medical Research, Parkville, Victoria, Australia; Department of Biochemistry and Molecular Biology, Biomedicine Discovery Institute, Monash University, Melbourne, VIC 3068, Australia; Department of Medical Biology, University of Melbourne, Melbourne, VIC 3068, Australia

**Author notes:** Contributed equally (Giang Thanh Nguyen-Dien, Brendan Townsend, Prajakta Kulkarni, Keri-Lyn Kozul).

## Abstract

Mitophagy must be carefully regulated to ensure that cells maintain appropriate numbers of functional mitochondria. The SCF^FBXL4^ ubiquitin ligase complex suppresses mitophagy by controlling the degradation of BNIP3 and NIX mitophagy receptors, and *FBXL4* mutations result in mitochondrial disease as a consequence of elevated mitophagy. Here, we reveal that the mitochondrial phosphatase PPTC7 is an essential cofactor for SCF^FBXL4^-mediated destruction of BNIP3 and NIX, suppressing both basal and induced mitophagy. Disruption of the phosphatase activity of PPTC7 is not required for BNIP3 and NIX turnover. Rather, a pool of PPTC7 on the mitochondrial outer membrane acts as an adaptor linking BNIP3 and NIX to FBXL4, facilitating the turnover of these mitophagy receptors. PPTC7 accumulates on the outer mitochondrial membrane in response to mitophagy induction or the absence of FBXL4, suggesting a homeostatic feedback mechanism that attenuates high levels of mitophagy. We mapped critical residues required for PPTC7-NIX/BNIP3 and PPTC7-FBXL4 interactions and their disruption interferes with both NIX/BNIP3 degradation and mitophagy suppression. Collectively, these findings delineate a complex regulatory mechanism that restricts NIX/BNIP3-induced mitophagy.

## Introduction

Cells eliminate excessive or damaged mitochondria through mitophagy, a selective form of autophagy^1,2^. The upregulation of mitophagy receptors BNIP3 and NIX on the mitochondrial outer membrane acts as a signal to recruit the autophagosome^3–5^, in conditions such as hypoxia^6–8^, and during the differentiation of specialized cell types like erythrocytes^9–11^, neurons^12,13^, cardiomyocytes^6,14^, keratinocytes^15^, and pro-inflammatory macrophages^13^. It has long been known that BNIP3 and NIX expression is acutely upregulated by transcription^16,17^, however, the molecular understanding of the mechanisms that restrict BNIP3 and NIX expression to prevent excessive mitophagy remains limited.

We and others have recently demonstrated that the abundance of BNIP3 and NIX receptors is regulated by the mitochondrially-localised SCF-FBXL4 E3 ubiquitin ligase complex^18–21^. FBXL4 is one of 69 F-box proteins that act as interchangeable substrate adaptors for SCF E3 ubiquitin ligase protein complexes. SCF^FBXL4^ localises to the mitochondrial outer membrane and mediates the constitutive ubiquitylation and degradation of BNIP3 and NIX to limit their abundance and thereby suppress mitophagy. The FBXL4 gene is mutated in MTDPS13 mtDNA depletion syndrome, a disease characterised by mitochondrial depletion caused by excessive mitophagy^22–24^.

Little is known regarding the upstream mechanisms that regulate BNIP3 and NIX mitophagy receptor recognition via FBXL4. Phosphorylation or other types of modifications could potentially either enhance or sterically preclude the recognition of BNIP3 and NIX by FBXL4. Alternatively, a cofactor might be required to regulate the assembly of the SCF complex or to bridge the interaction between FBXL4 with BNIP3 and NIX.

The PP2C phosphatase PPTC7 (Protein Phosphatase Targeting COQ7) localises to mitochondrial matrix ^25,26^. Despite this localization, it has been identified as an interactor of BNIP3 and NIX ^27–30^ and a suppressor of BNIP3 and NIX-dependent mitophagy ^31^. Intriguingly, PPTC7 knockout mice exhibit phenotypes reminiscent of FBXL4 KO mice with decreased mitochondria content, increased mitophagy and severe metabolic defects which are associated with perinatal lethality ^21,22,26,32^.

Here, we show that PPTC7 is a critical rate-limiting activator of FBXL4-mediated destruction of BNIP3 and NIX and is required to suppress excessive basal and mitophagy induced by pseudohypoxia. We propose that PPTC7 acts as an adaptor to enable the turnover of BNIP3 and NIX via the SCF^FBXL4^. An outer membrane form of PPTC7 interacts with BNIP3 and NIX as well as with FBXL4 and these interactions are required for BNIP3 and NIX turnover and mitophagy suppression. We functionally validate *in silico* predictions of the PPTC7-NIX and PPTC7-FBXL4 interactions to reveal critical residues required for the assembly of the PPTC7-NIX/BNIP3 and PPTC7-FBXL4, and consequently for the turnover of BNIP3 and NIX as well as mitophagy suppression. Together, these findings provide a molecular understanding of the mechanisms that restrict basal and NIX/BNIP3-stimulated mitophagy to prevent excessive mitophagy.

## Results

### PPTC7 and FBXL4 cooperate to mediate the turnover of BNIP3 and NIX

Corresponding with increased mitophagy, PPTC7 deficient cells and tissues have diminished steady-state levels of most mitochondrial proteins, except for BNIP3 and NIX, which instead exhibit a large increase in protein levels ^26,32^. To determine whether PPTC7 regulates BNIP3 and NIX expression at the level of protein stability, we generated PPTC7-deficient cell lines using CRISPR/Cas9-mediated gene disruption. Successful knockout was confirmed by immunoblotting using an anti-PPTC7 antibody, which detected bands migrating at 28 kDa, 32 kDa and 40 kDa, specifically in parental cell lines but not in PPTC7 KO lines (**Figure 1A, 1D, S1B-C**). Cycloheximide chase assays demonstrated that PPTC7 deficiency significantly increased the half-life of BNIP3 and NIX and resulted in the upregulation of BNIP3 and NIX at mitochondria, indicating that PPTC7 is required for BNIP3 and NIX turnover (**Figure 1A-C and S1A-B**). To assess whether PPTC7 impacts the transcriptional regulation of BNIP3 and NIX via HIF1α, we used the HIF1α inhibitor, echinomycin, to demonstrate that HIF1α inhibition did not abolish the accumulation of BNIP3 and NIX in PPTC7 KO cells (**Figure S1C**). The abnormally increased levels of BNIP3 and NIX in the PPTC7 KO lines were rescued upon re-expression of wild-type PPTC7 (**Figure 1C**). Collectively, our data suggests that PPTC7 is required for the turnover of BNIP3 and NIX.

**Figure 1.**
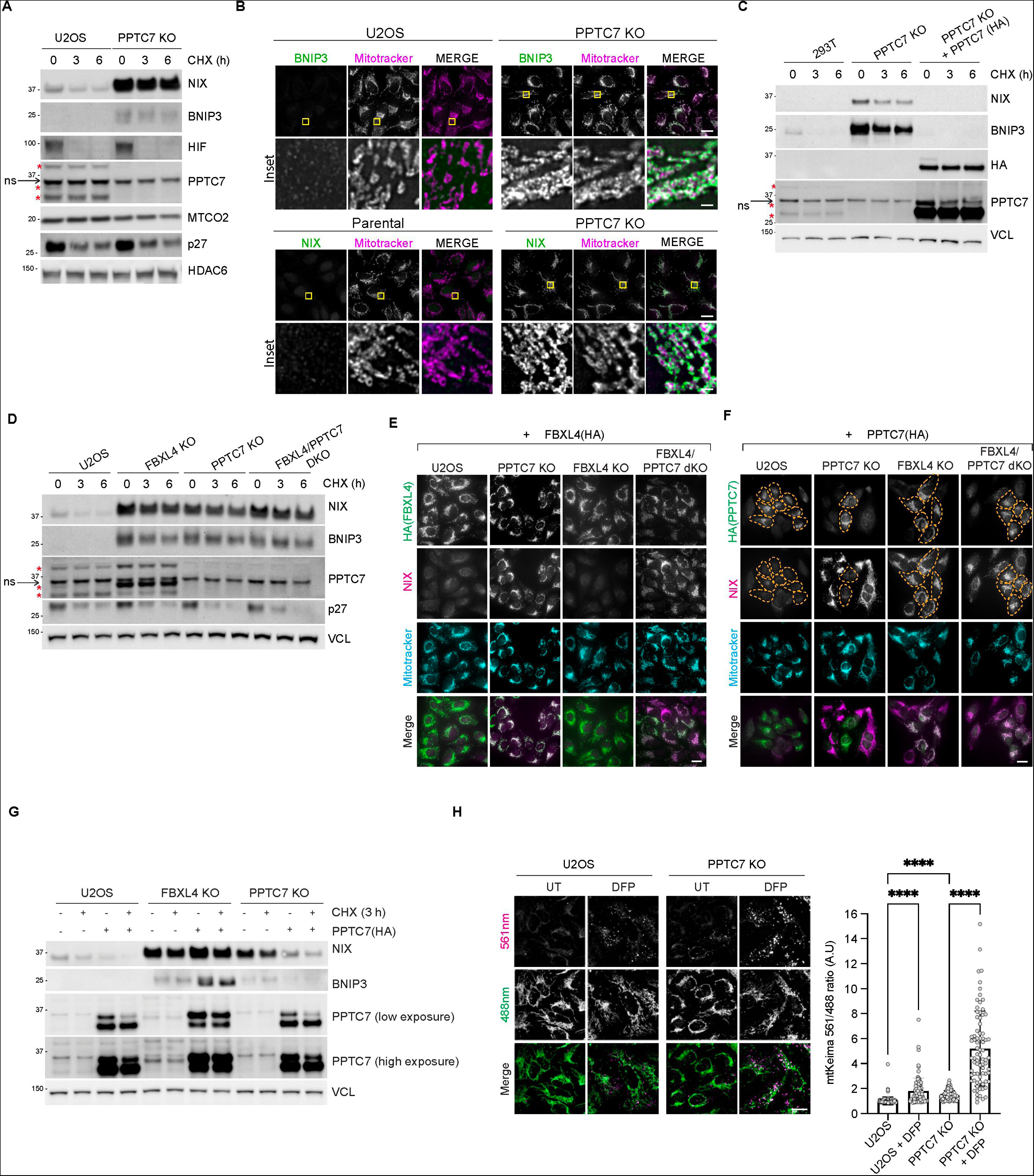
PPTC7 and FBXL4 coordinate the turnover of BNIP3 and NIX mitophagy receptors to suppress mitophagy. A. *BNIP3 and NIX are stabilized in PPTC7-deficient U2OS cells.* CRISPR-CAS9 was used to knockout PPTC7 in U2OS cells. Cells were treated with cycloheximide for the indicated times before immunoblotting as indicated. The red asterisks indicate PPTC7 specific bands at 28 kDa, 32 kDa, and 40 kDa. ns=non-specific band at ∼36 kDa. B. Immunofluorescence staining of BNIP3 and NIX demonstrating increased levels of both proteins at mitochondria in PPTC7 deficient cells U2OS cells. Cells were stained with BNIP3 or NIX (in green) and counterstained with MitoTracker (in magenta). Scale bar of main = 20 microns. Scale bar of inset = 1 micron. C. *Re-expression of PPTC7 into PPTC7-deficient cells reduces the stability of BNIP3 and NIX.* PPTC7-HA was transduced into PPTC7-deficient 293T cells. The half-lives of BNIP3 and NIX were analysed by immunoblotting after a cycloheximide chase. D. *Targeting PPTC7 and FBXL4 simultaneously does not further increase BNIP3 and NIX levels or stability.* PPTC7 was sequentially knocked out of previously generated FBXL4-deficient cells to make the double knockout of PPTC7 and FBXL4 (dKO FBXL4/PPTC7). The 32 kDa band of PPTC7 was upregulated in the FBXL4 KO cells. E. *FBXL4 requires PPTC7 to mediate the downregulation of NIX.* FBXL4-HA was transduced in parental, FBXL4 KO, PPTC7 KO, and FBXL4/PPTC7 dKO cells. NIX protein levels (magenta) were monitored in FBXL4(HA) expressing cells (green). Full figure in Appendix 1A. Scale bar = 20 microns. F. *PPTC7 requires FBXL4 to mediate the downregulation of NIX.* PPTC7-HA was expressed in parental, FBXL4 KO, PPTC7 KO, and FBXL4/PPTC7 dKO cells. NIX protein levels (magenta) were monitored in PPTC7(HA) expressing cells (green). To correlate NIX levels with PPTC7 expression, PPTC7 expressing cells (green) are outlined in orange dotted lines. Full figure in Appendix 1B. Scale bar = 20 microns. G. *FBXL4 is required for the PPTC7-mediated downregulation of BNIP3 and NIX.* PPTC7(HA) was transduced into U2OS, FBXL4 KO or PPTC7 KO cells and the half-lives of BNIP3 and NIX were monitored by immunoblotting. PPTC7(HA) rescue into PPTC7 KO cells causes a reduction of BNIP3 and NIX. In contrast, the expression of PPTC7(HA) into FBXL4 KO cells does not cause a reduction in either NIX or BNIP3. H. *PPTC7-deficiency leads to elevated mitophagy in basal conditions and after DFP treatment.* U2OS mtKeima PPTC7 KO cells were treated with DFP for 24 hours. Emission signals at neutral pH were obtained after excitation with the 458 nm laser (green), and emission signals at acidic pH were obtained after excitation with the 458 nm laser 561 nm laser (magenta). Mitophagy is represented as the ratio of mt-Keima 561 nm fluorescence intensity to mt-Keima 458 nm fluorescence intensity for individual cells normalised to the mean of the untreated condition. > 70 cells per condition were analysed. P values were derived from a non-parametric one-way ANOVA test. ****p<0.0001. Source data available. Scale bar = 20 microns.

The turnover of BNIP3 and NIX depends on the SCF^FBXL4^ ubiquitin ligase ^18,19,21^. To investigate whether PPTC7 and FBXL4 operate within the same or separate pathways to regulate BNIP3 and NIX turnover, we assessed the stability of BNIP3 and NIX in CRISPR knockout cell lines lacking either PPTC7, FBXL4, or both using a cycloheximide chase assay (**Figure 1D**). Our results demonstrated that the combined deficiency of both PPTC7 and FBXL4 did not lead to further upregulation of either BNIP3 or NIX compared with the individual knockout of PPTC7 or FBXL4, suggesting that FBXL4 and PPTC7 function in a shared pathway (**Figure 1D**). Strikingly, FBXL4 knockout cells displayed a notable increase of the 32 kDa (middle) form of PPTC7 (**Figure 1D**). The 32 kDa form of PPTC7 also accumulated after DFP treatment (**Figure S1C**).

Immunofluorescence-based complementation assays in FBXL4- and PPTC7-deficient cells showed that the ability of either PPTC7 or FBXL4 to mediate BNIP3 and NIX turnover depends on the presence of the other protein. Rescue of either FBXL4 or PPTC7 into their respective knockout cell lines resulted in the drastic downregulation of NIX levels compared with surrounding untransfected cells (**Figure 1E-F and S1E**). In contrast, FBXL4 overexpression was not able to mediate the downregulation of BNIP3 and NIX in the absence of PPTC7 (**Figure 1E and S1D**). Likewise, PPTC7 expression could not promote BNIP3 and NIX downregulation in the absence of FBXL4 (**Figure 1F-G and S1E**). Notably, in these experiments, PPTC7 deficiency did not affect the localization or levels of FBXL4 (**Figure 1F and S1D**). Thus, FBXL4 and PPTC7 require each other to mediate the downregulation of BNIP3 and NIX.

To determine whether PPTC7 levels are rate-limiting for BNIP3 and NIX turnover and consequently for mitophagy suppression, we examined the levels of BNIP3 and NIX in U2OS and 293T cells stably overexpressing PPTC7 (at approximately 100-fold for U2OS and 50-fold for 293T cells). BNIP3 and NIX levels were examined in steady-state conditions or after treatment with DFP, which is an iron chelator and HIF1α activator known to promote mitophagy via BNIP3 and NIX upregulation ^6,8^. The low levels of BNIP3 and NIX in steady-state conditions made it hard to detect a further decrease in levels upon PPTC7 overexpression in U2OS, however, we found that the overexpression of wild-type PPTC7 resulted in the downregulation of BNIP3 and NIX after DFP treatment in U2OS cells and 293T cells (**Figure S1F-H**). Consistent with its suppression of BNIP3 and NIX protein levels, the overexpression of PPTC7 suppressed DFP-induced mitophagy in U2OS cells (**Figure S1H**). Furthermore, PPTC7 knockout U2OS cells treated with DFP exhibited substantially more mitophagy than either condition alone, indicating that PPTC7 suppresses both basal and DFP-induced mitophagy (**Figure 1H**). Altogether, our experiments indicate that PPTC7 is rate-limiting for FBXL4-mediated BNIP3 and NIX turnover and the associated mitophagy suppression (**Figure 1H**).

### A population of PPTC7 localises to the outer mitochondrial membrane and interacts with FBXL4 and NIX/BNIP3

PPTC7 has been previously localized to the mitochondrial matrix ^25,33^, whereas BNIP3 and NIX are located at the mitochondrial outer membrane. We sought to clarify the sub-mitochondrial localization of PPTC7 to explore whether PPTC7 is located at both the outer mitochondrial membrane as well as inside the mitochondria. Like other mitochondrial proteins, PPTC7 possesses an N-terminal mitochondrial targeting pre-sequence (MTS) that is predicted to be proteolytically cleaved by mitochondrial proteases after import into the mitochondria ^34^, giving rise to a shorter processed form of the protein. Given the molecular weight difference predicted from the removal of the MTS, we posited that the different bands of PPTC7 might represent the precursor form of PPTC7 that has not been imported into mitochondria, and a processed shorter matrix form. To test this hypothesis, we conducted mitochondrial import assays, employing proteinase K to degrade proteins that reside on the outside of the mitochondria (i.e., that are not yet imported) (**Figure 2A**). We observed that the shorter molecular weight form of PPTC7 was resistant to proteinase K suggesting that it is inside the mitochondria. The 40 kDa (upper) form of PPTC7 was also proteinase K resistant, indicating that it likely resides in the matrix. Currently, the nature of the third 40 kDa form of PPTC7 remains unclear, however, we note that we only observe it for endogenous PPTC7 and not exogenous PPTC7.

**Figure 2.**
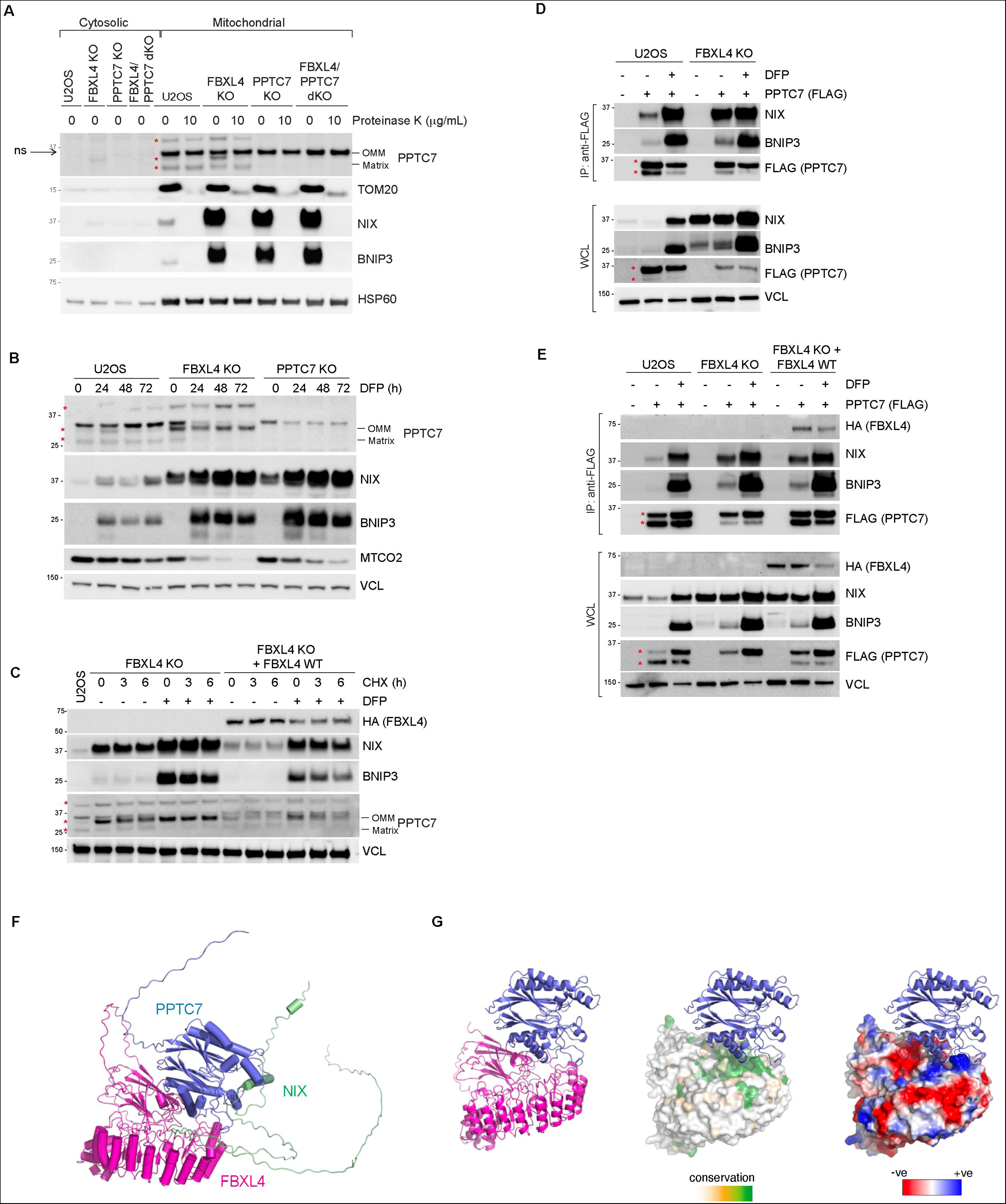
A population of PPTC7 localises to the mitochondria outer membrane to enable its interaction with BNIP3 and NIX and FBXL4. A. *The 32 kDa migrating form of PPTC7 is located on the mitochondrial outer membrane, while the lower migrating form of PPTC7 resides in the matrix.* Mitochondria were isolated from parental U2OS, FBXL4 KO, PPTC7 KO or PPTC7/FBXL4 dKO cells. The submitochondrial localisation of PPTC7 was determined by proteinase K treatment, which degrades proteins that are not imported within the mitochondria, i.e., outer membrane proteins. The red asterisks indicate PPTC7-specific bands. The arrow points to the non-specific band at 36 kDa which is enriched in mitochondria. B. *The outer membrane form of PPTC7 is induced by DFP treatment and in FBXL4 deficient cells.* U2OS cells or FBXL4 KO cells were treated with DFP for the indicated times. C. Analysis of the stability of the outer membrane form of PPTC7 in response to FBXL4 deficiency. FBXL4 KO cells and FBXL4 KO cells expressing FBXL4(HA) were subjected to cycloheximide chase assay in the presence or absence of DFP. DFP treatment and FBXL4 deficiency both extended the half-live of OM-PPTC7. D. *PPTC7 interacts with NIX/BNIP3 in basal conditions in an FBXL4-independent manner.* U2OS cells or FBXL4 KO cells were transfected with PPTC7-FLAG and treated with DFP for 24 hours. Cell lysates were immunoprecipitated with anti-FLAG beads, and the immuno-precipitates were analysed by immunoblotting as indicated. WCL = whole cell lysates. E. *PPTC7 interacts with FBXL4.* As in E, including FBXL4 KO cells rescued with HA-tagged FBXL4 WT. F. *AlphaFold2 multimer model of FBXL4, PPTC7 and NIX.* See also Figures S2F-G. G. The FBXL4 pocket is coloured to indicate surface sequence conservation or surface electrostatics using Consurf or Protein-Sol Patches server.

In contrast, we found that the 32 kDa (middle) version of PPTC7, which accumulates in FBXL4-deficient cells, is susceptible to proteinase K, indicating its localization on the mitochondrial outer membrane. Hereafter, we refer to the 32 kDa form of PPTC7 as outer membrane PPTC7 (OM-PPTC7), and the 28 kDa form of PPTC7 as inner mitochondrial PPTC7 (matrix-PPTC7).

We next explored the relative abundance of the outer/inner forms of PPTC7 in response to different mitophagy activators. In addition to FBXL4 deficiency, conditions known to upregulate BNIP3 and NIX-dependent mitophagy, such as the hypoxia-mimetics DFP and DMOG, resulted in the upregulation of the OM-PPTC7 (**Figure 2B-C and S2A-B**), largely correlating with BNIP3 and NIX levels. The greatest upregulation of OM-PPTC7 was observed in DFP-treated, FBXL4-deficient cells, and in this condition, the levels of matrix-PPTC7 were inversely correlated with OM-PPTC7. Note that the non-specific band detected by the PPTC7 antibody disappeared in the conditions associated with the highest mitophagy, which we presume is due to a decrease in mitochondrial content caused by elevated mitophagy. In the same conditions, the decrease in matrix-PPTC7 could theoretically reflect either compromised mitochondrial import of PPTC7 or high levels of mitochondrial degradation through mitophagy or a combination of both and warrants further investigation.

Next, we used affinity purification of FLAG-tagged PPTC7 to demonstrate that PPTC7 can robustly interact with both BNIP3 and NIX (**Figure 2E**) and FBXL4 (**Figure 2F**). The interaction between PPTC7 and NIX/BNIP3 was evident in basal conditions, as well as after DFP treatment (**Figure 2E-F**). We also established that FBXL4 is not required for the interaction between PPTC7 and NIX/BNIP3 using FBXL4-deficient cells (**Figure 2F**). In our experimental conditions, it was not possible to conclusively determine whether the interactions between PPTC7 with BNIP3, NIX or FBXL4 changed in response to DFP treatment since they were confounded by the changing levels of proteins after DFP (increased NIX/BNIP3 after DFP, and decreased FBXL4) and require further investigation. Lastly, we tested whether PPTC7 is required for the ability of FBXL4 to bind to SKP1 and CUL1, core members of the SCF complex. FBXL4 interacted equally with CUL1 and SKP1 in PPTC7 deficient cells, suggesting that PPTC7 is not required for SCF assembly (**Figure S2C**).

To confirm the direct interaction between PPTC7 and NIX/BNIP3 we tested the binding of BNIP3 and NIX synthetic peptide sequences to recombinant PPTC7 by isothermal titration calorimetry (ITC). We first tested the functionality of PPTC7 by measuring the association of the PPTC7 active site with divalent cations, observing binding of both Mg^2+^ and Mn^2+^, but with a much higher affinity for Mn^2+^ (**Figure S2E; Supplementary Table 1**) in line with the previous demonstration of Mn^2+^-dependent activity ^25^. Subsequently, we found that both BNIP3 and NIX peptides bound to PPTC7 directly, with similar although modest affinities (*K*_d_) of 20 and 37 μM respectively **(Figure S2F; Supplementary Table 2**).

Since BNIP3 and NIX demonstrated binding to PPTC7, but little to no binding with FBXL4 in our experimental conditions (**Figure S2B**), this led us to speculate that PPTC7 might serve as a bridge or scaffold between FBXL4 and NIX/BNIP3. To examine this at the molecular level, we modelled the interactions between PPTC7, FBXL4 and NIX/BNIP3 using AlphaFold2 (**Figure 2F and S2D-E**). Predictions of pairwise complexes or all three proteins together resulted in identical structural models. Alphafold2 predicts a high confidence interaction between FBXL4 and one surface of PPTC7, while a conserved sequence found in both BNIP3 and NIX associates with the active site of PPTC7. A conserved and negatively charged pocket formed between FBXL4’s N-terminal discoidin domain and C-terminal LRR domains surrounds PPTC7’s residues E125-K130 (**Figure 2F-G and S2G-H**). The binding of BNIP3 and NIX involves a highly extended peptide sequence of ∼25 residues including the distal end of BNIP3/NIX’s non-canonical BH3 domains, centred on a highly conserved sequence, which we hereafter refer to as the SRPE sequence (encompassing residues 187-190 and 146-149 and in BNIP3 and NIX respectively). The conserved Ser sidechain is predicted to bind the catalytic pocket, precisely where a phosphorylated substrate would be expected to interact. These structural predictions are consistent with PPTC7 serving to position NIX/BNIP3 to be substrates of the SCF^FBXL4^ E3 complex.

### Disruption of the catalytic activity of PPTC7 does not affect BNIP3 and NIX turnover in basal conditions

BNIP3 and NIX are known to be phosphoproteins, and PPTC7 knockout systems have shown elevated phosphorylation at specific residues on BNIP3 and NIX^32^, albeit not within the regions of BNIP3 and NIX predicted to interact with PPTC7. Specifically, interrogation of existing phosphorylation databases finds no evidence of phosphorylation on NIX’s Ser146 or BNIP3’s Ser187, which are situated within the PPTC7 active site. Furthermore, phospho-proteomic analysis of affinity-purified BNIP3 from PPTC7 KO cells did not detect phosphorylation at Ser187 (**Supplementary Table 3**).

To determine whether the phosphatase activity of PPTC7 is required for BNIP3 and NIX turnover, we engineered mutant versions of PPTC7 predicted to have disrupted phosphatase activity. This involved mutating the active site residues, which were identified through a comparison of the AlphaFold2 prediction of PPTC7 (ID: AF-Q8NI37-F1) with the crystal structure of the similar PPM/PP2C family homologue photosystem II (PSII) core phosphatase (PBCP) (PDB ID: 6AE9)^35^. Specifically, we substituted the metal-binding aspartate residues at positions Asp78, Asp233, and Asp290 in PPTC7 (**Figure 3A and S3A**). We found that PPTC7 mutants in which Asp78 and Asp223 were mutated to alanine (PPTC7-D78A and PPTC7-D223A) were unable to rescue BNIP3 and NIX turnover when complemented into PPTC7-deficient cells (**Figure S3B-C**). However, these mutations also significantly reduced the binding of PPTC7 to BNIP3 and NIX (**Figure S3D**), either due to the loss of metal-binding required to coordinate the Ser sidechain in the SRPE NIX/BNIP3 binding sequences, or possibly due to conformational rearrangements. Thus, it was not possible to ascertain if the lack of turnover was due to a lack of phosphatase activity or reduced binding to NIX/BNIP3.

**Figure 3.**
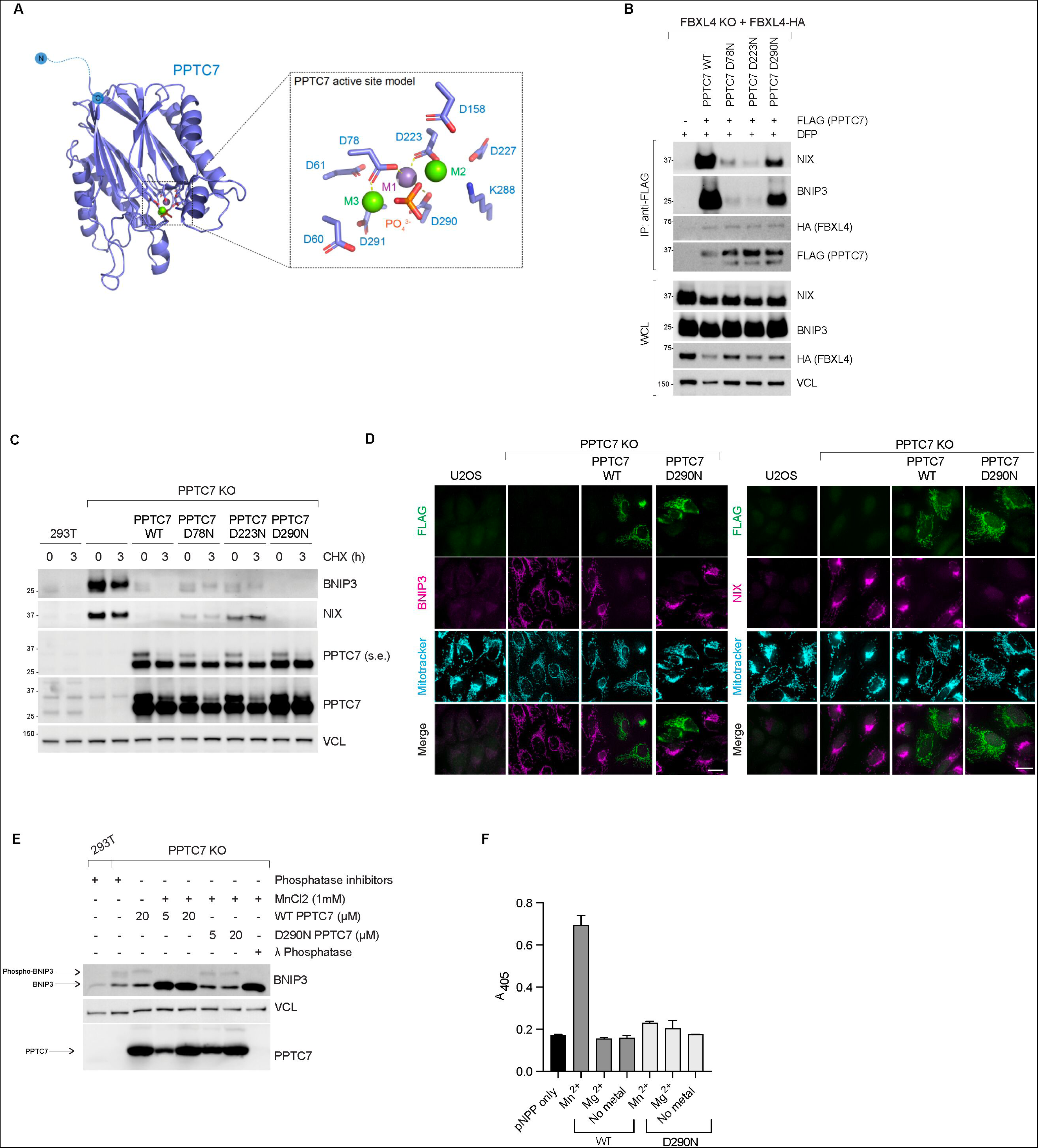
Disruption of the catalytic activity of PPTC7 does not affect BNIP3 and NIX turnover in basal conditions. A. *AlphaFold2 model of PPTC7 with active site residues indicated.* This is based on a comparison to the PBCP structure (PDB ID 6AE9). The three putative metal ions and the incoming phosphate group are modelled based on an alignment of the PPTC7 and PBCP structures. B. *Binding to BNIP3 and NIX is partially preserved in the PPTC7-D78N and PPTC7-D290N mutants.* FBXL4 KO complemented with FBXL4(HA) were transfected with PPTC7-FLAG or D78N, D223N, and D290N mutants. Cells were treated with DFP for 24 hours to visualise binding better. Cell lysates were immunoprecipitated with anti-FLAG beads, and the immuno-precipitates were analysed by immunoblotting as shown. C. *PPTC7-wildtype can dephosphorylate BNIP3 but PPTC7-D290N cannot.* Electrophoretic mobility shift assays of BNIP3 using phos-tag gels were performed to assess the phosphorylation status of BNIP3. Purified PPTC7 or PPTC7-D290N were incubated with 293T PPTC7 KO cell lysates. Wild-type PPTC7 was able to dephosphorylate BNIP3 in a Mn2+-dependent manner as demonstrated by the loss of the upper form of BNIP3. However, PPTC7-D290N was not able to dephosphorylate BNIP3 in the same conditions. Lamda-phosphatase is used as a control to demonstrate that the upper band of BNIP3 seen on the phos-tag gels is the phosphorylated species. D. *In vitro pNPP dephosphorylation assay using purified PPTC7-wildtype and PPTC7-D290N.* The dephosphorylation of pNPP was measured at 405 nm after a 15 minute reaction. 10 mM pNPP was added to 4 µM PPTC7 in the presence or absence of either MnCl_2_ or MgCl_2_. E. *PPTC7-D290N can restore the turnover of BNIP3 and NIX to a similar extent as wild-type PPTC7.* The PPTC7 KO cells were rescued with wild-type PPTC7-FLAG or PPTC7-D78N, PTPC7-D223N or PPTC7-D290N variants. Cells were treated with cycloheximide for 3 hours before harvesting. Samples were lyzed, and immunoblotting was performed. F. *PPTC7-D290N can rescue the degradation of BNIP3 and NIX to a similar extent as wild-type PPTC7.* NIX and BNIP3 protein levels (magenta) were monitored in PPTC7(FLAG) expressing cells. To correlate NIX/BNIP3 levels with PPTC7 expression, PPTC7 expressing cells (green) are outlined in orange dotted lines.

To preserve the metal binding activity of PPTC7 and thus the interaction between PPTC7 and NIX/BNIP3, we next substituted the polar aspartate residues with alternate polar asparagine residues (D78N, D223N, or D290N). This change preserved partial binding between PPTC7 and NIX/BNIP3, with the PPTC7-D290N variant displaying the greatest binding compared with wild-type PPTC7 (**Figure 3B**).

To determine if the catalytic activity of PPTC7 is important for BNIP3 and NIX degradation, we expressed PPTC7-wildtype or PPTC7-aspartate to asparagine active site mutants in PPTC7 KO cells. Complementation assays demonstrated that PPTC7-D290N can fully rescue the turnover of BNIP3 and NIX as well as wild-type PPTC7 (**Figure 3C-D**), with partial rescue observed for PPTC7-D78N and PPTC7-D223N. We confirmed that PPTC7-D290N had compromised catalytic activity compared with PPTC7-wildtype by assessing the BNIP3 phospho-migration shift, finding purified wild-type PPTC7 could dephosphorylate BNIP3 however PPTC7-D290N could not (**Figure 3E**). PPTC7-D290N was also confirmed to be catalytically defective using pNPP phosphatase assays (**Figure 3F**). Our data demonstrates that although BNIP3 (and likely NIX) can be dephosphorylated by PPTC7 (**Figure 3E**), the FBXL4-mediated degradation of BNIP3 and NIX is still mediated by the catalytically defective PPTC7, suggesting that the turnover of PPTC7 depends on its presence, but not its full activity.

### NIX/BNIP3-PPTC7 interactions are critical for NIX/BNIP3 turnover and mitophagy suppression

To further test the significance of the interaction between NIX/BNIP3 and PPTC7, we sought to disrupt the interaction between NIX/BNIP3 and PPTC7. Based on our models of the PPTC7 complex with either BNIP3 or NIX using Alphafold2 ^36^, Tyr179 and Asn181 in PPTC7 were predicted to be critical residues required for binding of NIX/BNIP3 without affecting the PPTC7 active site (**Figure 3A**). On the NIX/BNIP3 interface, Trp144 in the BH3 domain, and a highly conserved SRPE region in NIX/BNIP3 were predicted to interact with PPTC7’s active site (residues 187-190 in BNIP3 and 146-149 and NIX) (**Figure 3A and 3D**).

To test the Alphafold2 prediction, we made the following mutations in PPTC7: Y179D to disrupt the interface with NIX/BNIP3 and N181E to disrupt the local pocket around the phosphoserine-binding catalytic site (**Figure 4A**). First, we tested whether these mutants lose binding to NIX/BNIP3 using an anti-FLAG affinity pulldown assay. Our results demonstrated that PPTC7-Y179D and PPTC7-N181E have reduced binding to both BNIP3 and NIX (**Figure 4B**), indicating that Tyr179 and Asn181 residues are important for the interaction with NIX/BNIP3. In contrast, Gln128 and Lys130 lie in the putative FBXL4 binding site, and Q128R and K130E mutations in PPTC7 did not affect NIX/BNIP3 binding, as predicted (**Figure 4B**), acting as controls for testing NIX/BNIP3 interactions.

**Figure 4.**
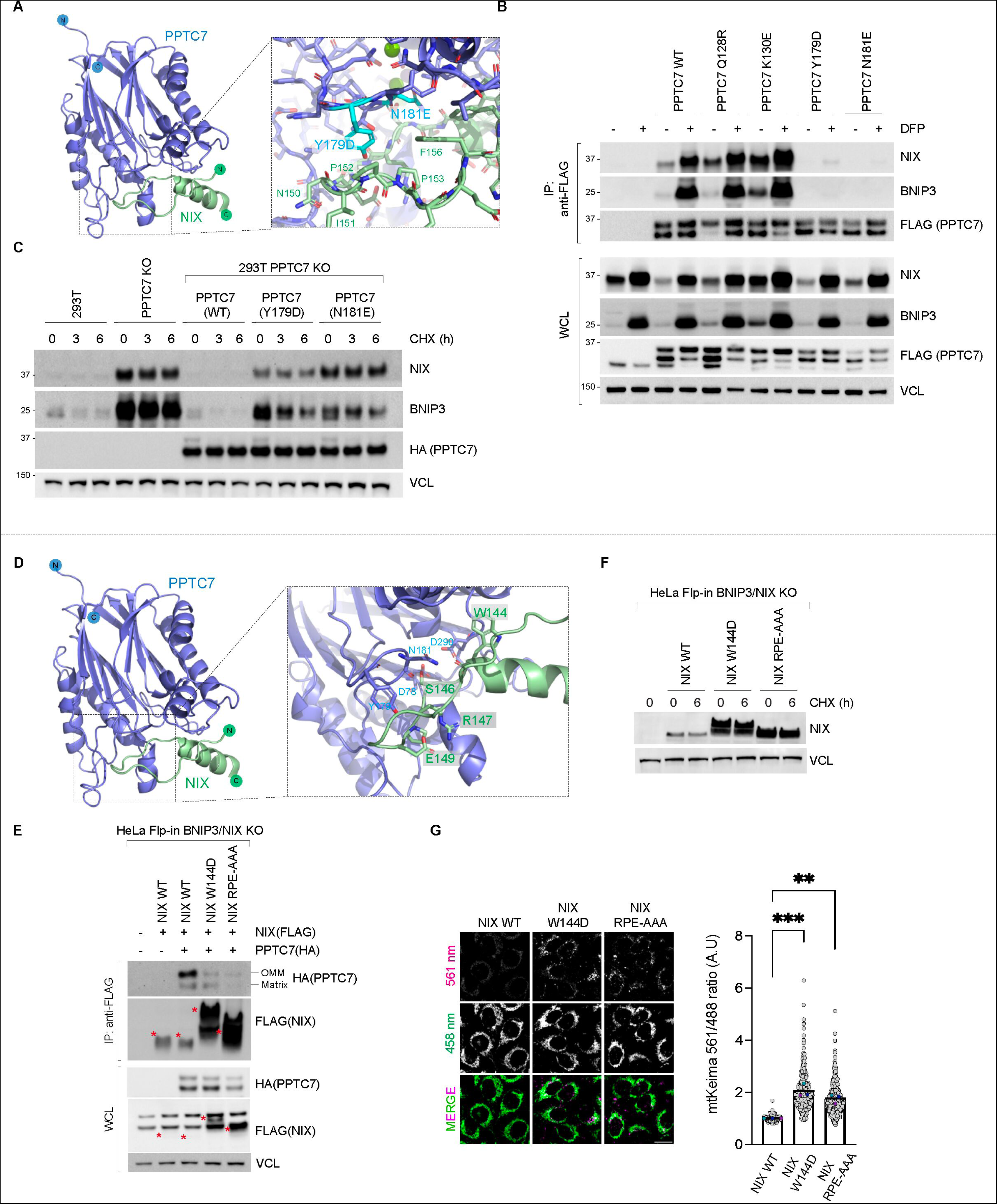
The NIX-PPTC7 interaction is critical for NIX turnover and mitophagy suppression. A. *AlphaFold2 prediction of the PPTC7-NIX complex.* Key residues in the PPTC7 interface are highlighted in cyan. B. *PPTC7-Y179D and PPTC7-N181E variants are unable to interact with BNIP3 and NIX*. PPTC7(FLAG) and specified mutants were transfected into U2OS cells. Cells were treated with DFP for 24 hours. Cell lysates were precipitated with anti-FLAG affinity resin and the immuno-precipitates were analysed by immunoblotting. C. *PPTC7-Y179D and PPTC7-N181E variants are unable to mediate the turnover of BNIP3 and NIX.* PPTC7(HA)-WT, PPTC7-Y179D or PPTC7-N181E were expressed in PPTC7 KO 293T cells. BNIP3 and NIX stability was assessed using a cycloheximide chase. D. *AlphaFold2 prediction of the PPTC7-NIX complex.* Key residues in the NIX interface including Arg147 are highlighted. E. *Residues W144 and RPE_147-149_ in NIX are critical for binding to OM-PPTC7.* PPTC7(HA) was transduced into HeLa Flp-in NIX/BNIP3 double KO cells stably expressing inducible FLAG-tagged NIX-WT or NIX mutants. NIX expression was induced with doxycycline for 24 hours. Cell lysates were immunoprecipitated with anti-FLAG beads, and the immuno-precipitates were analysed by immunoblotting. The levels of NIX-W144D and NIX-RPE/AAA were significantly higher than NIX-wild-type. Red asterisks mark the NIX(FLAG) proteins. The different migration of the point mutants by electrophoresis could be due to the change in charge of the residues or because PPTC7 is unable to dephosphorylate NIX. F. *NIX-W144D and NIX-RPE/AAA are stabilized in comparison to wild-type NIX.* HeLa Flp-in NIX/BNIP3 double KO cells expressing FLAG-tagged NIX WT or NIX mutants were subjected to a cycloheximide chase. G. *Expression of hyper-stable NIX-W144D and NIX-RPE/AAA leads to an increase in basal levels of mitophagy.* Hela Flp-In NIX knockout/ Hela Flp-In NIX/BNIP3 double knockout Keima cells stably expressing NIX and NIX mutants were treated with doxycycline for 48 hours and mitophagy was evaluated using live-cell confocal fluorescence microscopy. Mitophagy is represented as the ratio of mt-Keima 561 nm fluorescence intensity divided by mt-Keima 458 nm fluorescence intensity for individual cells normalised to the untreated U2OS cells. Translucent grey dots represent measurements from individual cells. Colored circles represent the mean ratio from independent experiments. The centre lines and bars represent the mean of the independent replicates +/- standard deviation. 4 data points are outside the axis limits. P values were calculated based on the mean values using a one-way ANOVA (*P<0.05, **P<0.005, ***P<0.001, ****P<0.0001). N=3.

Secondly, we tested whether the disruption of PPTC7’s ability to interact with NIX/BNIP3 disrupted the ability of PPTC7 to mediate NIX/BNIP3 turnover. As hypothesised, the loss of PPTC7’s interaction with BNIP3 and NIX resulted in a loss of its ability to mediate BNIP3 and NIX turnover (**Figure 4C**). We performed rescue assays in PPTC7 KO cell lines using either wild-type PPTC7 or the variants in PPTC7 with defective binding to NIX/BNIP3: PPTC7-Y179D and PPTC7-N181E. We found that, unlike wild-type PPTC7 which reduced the elevated BNIP3 and NIX levels, PPTC7-Y179D and PPTC7-N181E displayed elevated NIX/BNIP3 compared with wild-type PTC7 as assessed using a cycloheximide chase assay (**Figure 4C-D**). The mutants localised to the mitochondria as expected (**Figure 4D**).

To functionally validate the regions on BNIP3 and NIX required for their interaction with PPTC7, a series of NIX or BNIP3 mutation constructs were expressed in NIX/BNIP3 double knockout cells: NIX-W144D, NIX-Δ140-150, NIX-R147D, and triple substitutions of R-P-E_(147-149)_ to either alanines (RPE-AAA) or aspartic acid and alanines (RPE-DAA). We performed FLAG affinity purification assays in HeLa Flp-In NIX/BNIP3 double KO cells co-expressing doxycycline-inducible FLAG-tagged wild-type NIX or NIX mutants along with HA-tagged PPTC7 (**Figure 4E and S4C**). Indeed, we found that despite the lower levels of wild-type NIX, only wild-type NIX was able to bind to PPTC7 (**Figure 4E and S4C**), whereas NIX-W144D, NIX-RPE/AAA, NIX-RPE/DAA and NIX--Δ140-150 were not. Fitting with the hypothesis that the interaction occurs at the outer membrane, the OM-form of PPTC7 bound preferentially to NIX. The expression of PPTC7-WT promoted the downregulation of wild-type NIX, but not the PPTC7-binding NIX variants, supporting the model that the binding of PPTC7 to NIX is required for its degradation. Therefore, the region surrounding the SRPE domain in NIX is required for functional interaction with PPTC7.

Having demonstrated that NIX’s SRPE and nearby residues are important for PPTC7 binding, we next assessed their stability. Supporting the Alphafold2 model, each of the NIX mutants displayed higher expression and longer half-lives than wild-type NIX (**Figure 4F and S4D**). These results suggest that the PPTC7-binding region in NIX plays an important role in its turnover.

Since elevated levels of BNIP3 and NIX correlate with increased mitophagy, we tested the hypothesis that the expression of the PPTC7-resistant hyper-stable NIX mutants would result in elevated mitophagy using a mtKeima assay (**Figure 4H and S4D**). In our conditions, the expression of wild-type NIX does not induce mitophagy since it is expressed at low levels, enabling a direct comparison of how stabilizing NIX by preventing its interaction with PPTC7 influences mitophagy. We expressed NIX-RPE/AAA, NIX-RPE/DAA and NIX-W144D in mt-Keima-expressing and compared the induction of mitophagy to that induced by wild-type NIX (**Figure 4G and S4E**). We found that the expression of NIX-RPE/AAA and RPE/DAA resulted in substantially elevated mitophagy in cells compared with wildtype NIX, indicating that the stabilisation of NIX due to loss of its binding to PPTC7 results in hyperactivation of mitophagy.

Taken together, these results validate the Alphafold2 modelling and demonstrate the importance of the BH3-SRPE region of NIX/BNIP3 and the Tyr179 and Asn181 residues of PPTC7 in NIX-PPTC7 binding. Furthermore, the results demonstrate that PPTC7 interaction is important for NIX/BNIP3 turnover and mitophagy suppression.

### The FBXL4-PPTC7 interaction is critical for mitophagy receptor turnover and mitophagy suppression

To test whether PPTC7’s interaction with FBXL4 is required for BNIP3 and NIX turnover, we proceeded to explore PPTC7’s interaction with FBXL4 using AlphaFold2 modelling. The predictions indicated that FBXL4’s Met71 and Arg544 are important for interaction with PPTC7 (**Figure 5A**), therefore we generated the following FBXL4 mutants to test binding to PPTC7: FBXL4-M71E, FBXL4-R544E or FBXL4-M71E/R544E. Consistent with the structural models, we found that FBXL4 mutants displayed weaker binding to PPTC7 than wild-type FBXL4, with the greatest reduction in binding to PPTC7 observed for FBXL4-M71E/R544E (**Figure 5B**). The binding of BNIP3 and NIX to PTTC7 was greatly increased in cells expressing the FBXL4 mutants that were unable to bind to PPTC7 (FBXL4-M71E, M544E, and M71E/R544E). This increase in binding is likely due to their increased levels, rather than any increase in affinity although our data does not rule this out. However, the result confirms that the interaction between FBXL4 and PPTC7 is not required for the interaction between PPTC7 and NIX/BNIP3 (**Figure 5B**).

**Figure 5.**
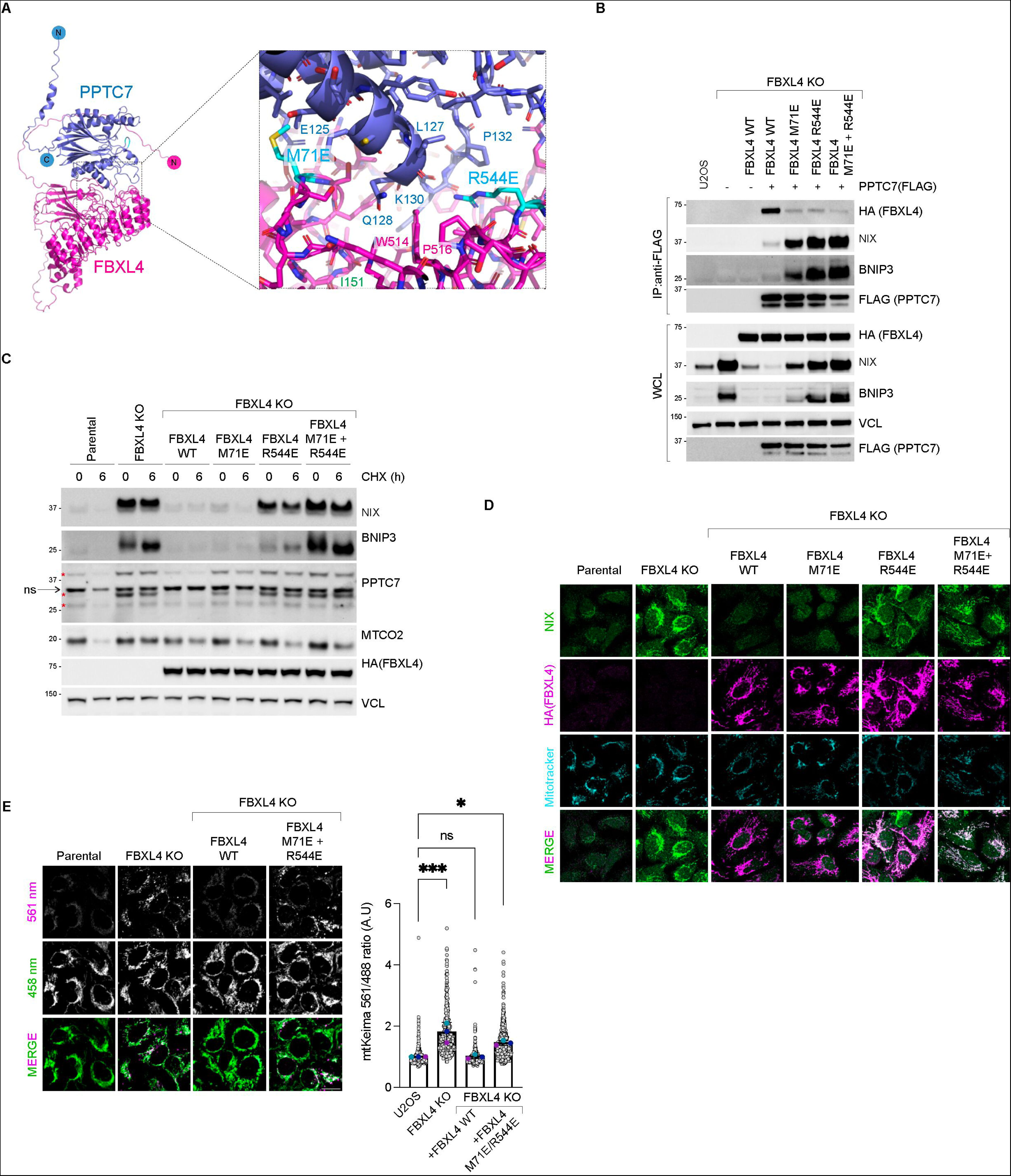
The FBXL4-PPTC7 interaction is required for BNIP3 and NIX turnover and mitophagy suppression. A. *Alphafold2 structural modelling of FBXL4 in complex with PPTC7.* Alphafold2 predicts a high confidence interaction between FBXL4 and PPTC7 centred on Met71 and Arg544 in FBXL4. B. *Met71 and Arg544 in FBXL4 are involved in the interaction with PPTC7.* FBXL4 knockout cells were complemented with HA-tagged wild-type FBXL4, FBXL4-M71E, FBXL4-R544E or FBXL4-M71E/R544E. Cells were transfected with FLAG-tagged PPTC7 as indicated, lysed, and subjected to affinity purification using anti-FLAG resin. WCL = whole cell lysates. C. *FBXL4-M71E, FBXL4-R544E or FBXL4-M71E/R544E variants are unable to mediate BNIP3 and NIX downregulation and destabilisation.* U2OS FBXL4 KO cells were rescued with wild-type FBXL4(HA) or specified variants. D. *Localization of FBXL4-M71E, FBXL4-R544E or FBXL4-M71E/R544E variants.* U2OS FBXL4 KO cells expressing FBXL4(HA) wildtype or specified variants were fixed and stained for HA to detect FBXL4 (in magenta) or NIX (green). NIX levels are inversely correlated with the ability of FBXL4 to bind to PPTC7. E. *FBXL4-M71E/R544E is less efficient than wild-type FBXL4 in mediating mitophagy suppression.* U2OS mt-Keima cells, U2OS mt-Keima FBXL4 KO cells and FBXL4 KO cells rescued with FBXL4 constructs were analyzed by confocal microscopy to quantify mitophagy.

**Figure 6.**
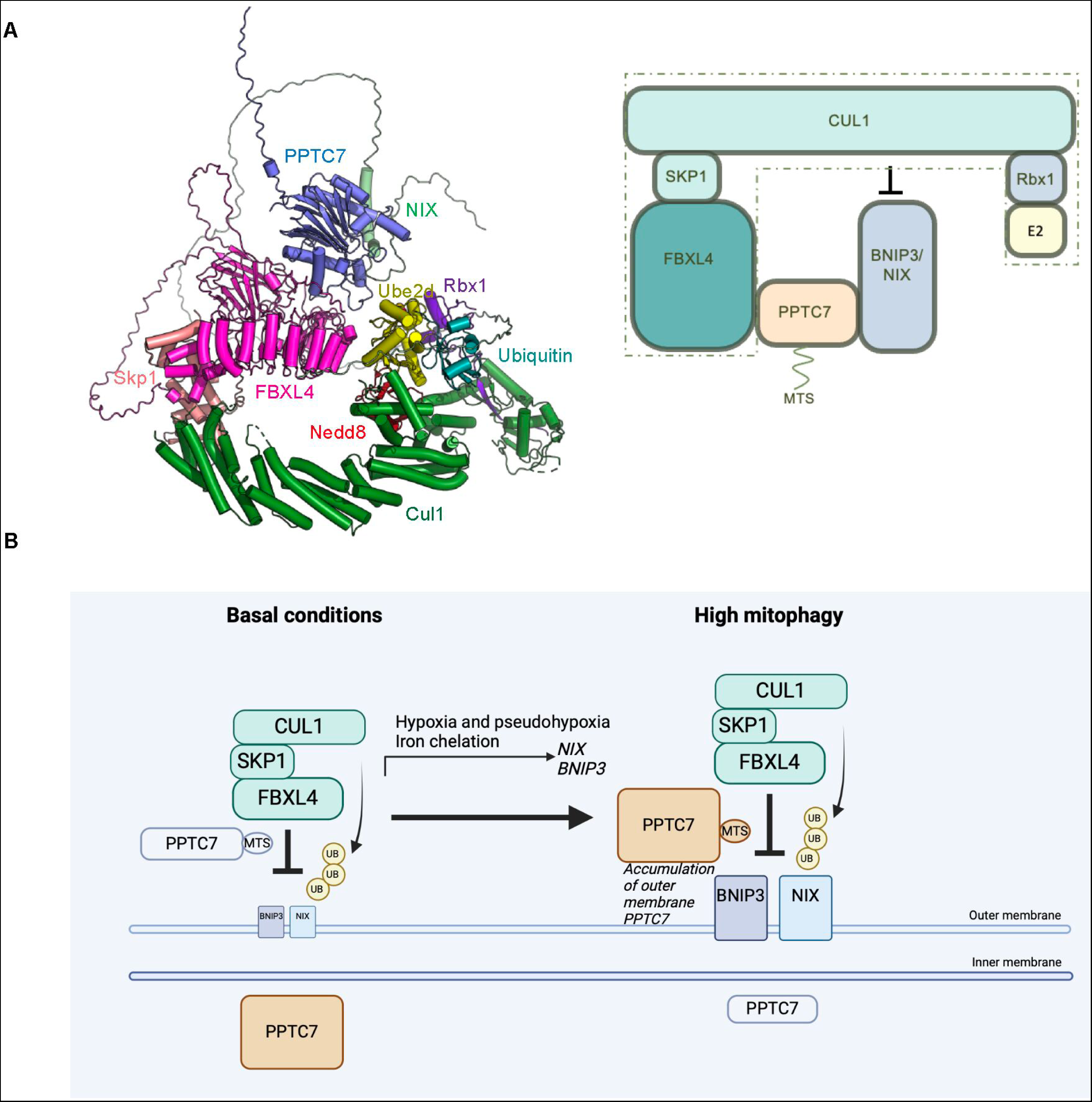
Working model for FBXL4 and PPTC7 mediated turnover of BNIP3 and NIX for mitophagy suppression. A. Model of the combination of Alphafold2 PPTC7-BNIP3-FBXL4 with the structure of Skp1-Cul1-Ube2d-Ub-Nedd8 (Beok 2020, PDB ID 6TTU). PPTC7 interacts with BNIP3/NIX and with FBXL4. One interpretation of our data is that PPTC7 bridges the interaction between FBXL4 and BNIP3/NIX to position BNIP3/NIX substrates for productive ubiquitylation by SCF-FBXL4. B. *Working model for the role of PPTC7 in mitophagy suppression.* In steady-state conditions, low levels of PPTC7 localize at the mitochondrial outer membrane to mediate the constitutive turnover of BNIP3/NIX, and the majority of PPTC7 remains in the mitochondrial matrix. PPTC7 accumulates at the mitochondria outer membrane in conditions of high mitophagy such as pseudohypoxia to dampen mitophagy. PPTC7 accumulation on the outer membrane may be a result of active retention mechanisms or defective mitochondrial import.

We next assessed the functional significance of this interface by assessing whether the loss of the binding between FBXL4 and PPTC7 affects the ability of FBXL4 to mediate the turnover of BNIP3 and NIX. Rescue assays were performed in FBXL4 KO cells expressing FBXL4 wild-type, FBXL4-M71E, FBXL4-E544E, or FBXL4-M71E/R544E and the stability of BNIP3 and NIX were assessed using a cycloheximide chase assay (**Figure 5C**). We found that the double mutant of FBXL4-M71E/R544E was unable to reduce the stability of BNIP3 and NIX to the same level as FBXL4-wild-type. Similar to FBXL4 deficiency, the outer-membrane form of PPTC7 was also stabilized in cells expressing FBXL4 variants unable to bind to PPTC7. Notably, despite their reduced ability to downregulate BNIP3 and NIX (**Figure 5B-C**), the FBXL4 variants localized, like wild-type FBXL4, to mitochondria (**Figure 5D**).

Finally, we validated that these residues are important for the ability of FBXL4 to suppress mitophagy, finding that FBXL4-M71E/R544E was also less effective at suppressing mitophagy when reconstituted into FBXL4-deficient cells, correlating with increased BNIP3 and NIX levels in cells expressing the variants compared with wild-type FBXL4 (**Figure 5E**). In all, these results suggest that the interaction with PPTC7 is required for FBXL4 to regulate NIX/BNIP3 turnover and for the ability of FBXL4 to suppress mitophagy.

## Discussion

In this study, we provide evidence that PPTC7 facilitates the SCF^FBXL4^-mediated turnover of BNIP3 and NIX. Our findings suggest that PPTC7 is a critical rate-limiting factor determining the amount of BNIP3 and NIX turnover and that it operates directly at the outer mitochondrial membrane by interacting with NIX/BNIP3 as well as with FBXL4, rather than employing an indirect mechanism.

PPTC7 accumulates at the mitochondrial outer membrane in response to pseudohypoxia and other conditions associated with elevated BNIP3/NIX-mediated mitophagy. The observed increase in outer-membrane PPTC7 in response to high mitophagy suggests a potential homeostatic feedback mechanism to limit excessive mitophagy. However, it remains incompletely understood how PPTC7’s localisation to the outer membrane is regulated. PPTC7 may be co-regulated with BNIP3 and NIX by FBXL4-mediated turnover. Alternatively, homeostasis may be achieved by modulating the rate of PPTC7 import into mitochondria. The accumulation of outer membrane PPTC7 may be a consequence of defective protein import, active retention mechanisms, or a combination of these possibilities. It also remains an open question whether this regulation might occur in a localized manner, perhaps targeting specific mitochondria.

While we were preparing this manuscript, a study by Sun and colleagues was published, which contains highly complementary findings to ours about the regulation of BNIP3 and NIX through the FBXL4 and PPTC7^37^. Their findings, akin to ours, show that PPTC7 acts as a critical and limiting factor in governing the FBXL4-mediated degradation of BNIP3 and NIX. Additionally, both studies highlight the dual localisation of PPTC7 to the matrix as well as the mitochondria outer membrane to enable its interaction with BNIP3 and NIX. Similarly, using different approaches to inhibit PPTC7, Sun et al suggest that the full catalytic activity of PPTC7 is not required for the turnover of BNIP3 and NIX, paralleling our observations. Sun et al additionally present elegant physiological data suggesting that the upregulation of PPTC7 in the context of liver during fasting conditions is required to maintain mitochondrial numbers in this organ. An interesting area needing further clarification in the future is the working model for how PPTC7 scaffolds the essential interactions between FBXL4 and NIX/BNIP3 and/or the SCF complex itself. In this regard, our data diverges from the working model of Sun *et al*, suggesting that rather than promoting SCF complex assembly through CUL1 recruitment, PPTC7 acts as an adaptor between FBXL4 and BNIP3. Further investigation incorporating structural and mechanistic data is required to reconcile these interesting findings. Altogether, the considerable overlap between our findings and those of Sun and colleagues underscores the indispensable role of the PPTC7-FBXL4 axis in suppressing mitophagy.

In silico modelling suggests that PPTC7 bridges the interaction of BNIP3 and NIX with FBXL4, enabling the effective positioning of the SCF complex for productive ubiquitylation of BNIP3/ NIX. This conclusion is further supported by functional assays, which demonstrate that disrupting the interactions between NIX and PPTC7, as well as between FBXL4 and PPTC7, results in the stabilization of BNIP3 and NIX, leading to increased basal mitophagy. Our results suggest that PPTC7 binds to directly to substrates BNIP3 and NIX, rather than the SCF components. The PPTC7 scaffold function we propose resembles the function of the CKS1 accessory factor, which plays a critical role in facilitating the interaction between FBXL1 (also known as SKP2) and its substrate, the cyclin-dependent kinase (Cdk) inhibitor p27^38^.

We note that we have not yet investigated whether certain conditions modulate FBXL4-PPTC7-NIX/BNIP3 interactions. These could be local signalling events, like oxidative stress, or global conditions like starvation. Moreover, although our structure-function analyses support the significance of individual structural interfaces, it remains possible that PPTC7 may interact in a mutually exclusive manner with either NIX/BNIP3 or FBXL4 since we have not provided experimental evidence for the existence of a trimeric complex.

Future research should address the importance of PPTC7’s catalytic activity in BNIP3 and NIX degradation. Phosphorylation of BNIP3 and NIX has been shown to influence their stability and/or their capacity to promote mitophagy ^5,39,40^ ^41,42^, suggesting that dephosphorylation may be required for their turnover and/or mitophagy suppression. Although our data suggest that PPTC7’s catalytic activity is largely dispensable for BNIP3 and NIX degradation, it remains possible that small amounts of catalytic activity from the PPTC7-D290N mutant are sufficient for turnover of BNIP3 and NIX in basal conditions. Alternatively, the catalytic activity of PPTC7 may be more important in certain conditions with elevated BNIP3 and NIX, such as during hypoxia or iron chelation. PPTC7-mediated dephosphorylation may promote critical interactions required for FBXL4 to mediate BNIP3 and NIX turnover. For instance, another F-box protein, FBXL2, interacts with the PTPL1 phosphatase, which dephosphorylates p85β on Tyr-655, thereby promoting p85β binding to FBXL2 and subsequent degradation ^43^. Both BNIP3 and NIX contain a serine residue (Serine146 in NIX and Ser187 in BNIP3) that directly interacts with the active site of PPTC7, where a phosphorylated substrate would typically bind. However, it is important to note that we have not yet found evidence for phosphorylation of this serine residue in existing literature or our phosphor-proteomic studies. Lastly, whether the catalytic activity or levels of PPTC7 and/or SCF-FBXL4 are regulated in response to environmental conditions remains a topic for further exploration.

On the other hand, if the enzymatic activity of PPTC7 is indeed dispensable for its regulation of NIX and BNIP3 turnover, this raises the question of the actual role of its catalytic activity. It is possible that PPTC7-dependent dephosphorylation of BNIP3 and NIX, or FBXL4—or perhaps regulatory proteins such as mitochondrial import proteins^26^—could serve a distinct role on the outer membrane to suppress mitophagy. Gaining insights into the kinases that counteract PPTC7’s function will be crucial for understanding how it contributes to the homeostatic regulation of mitophagy.

## Figure Legends

**Supplementary Figure 1.**
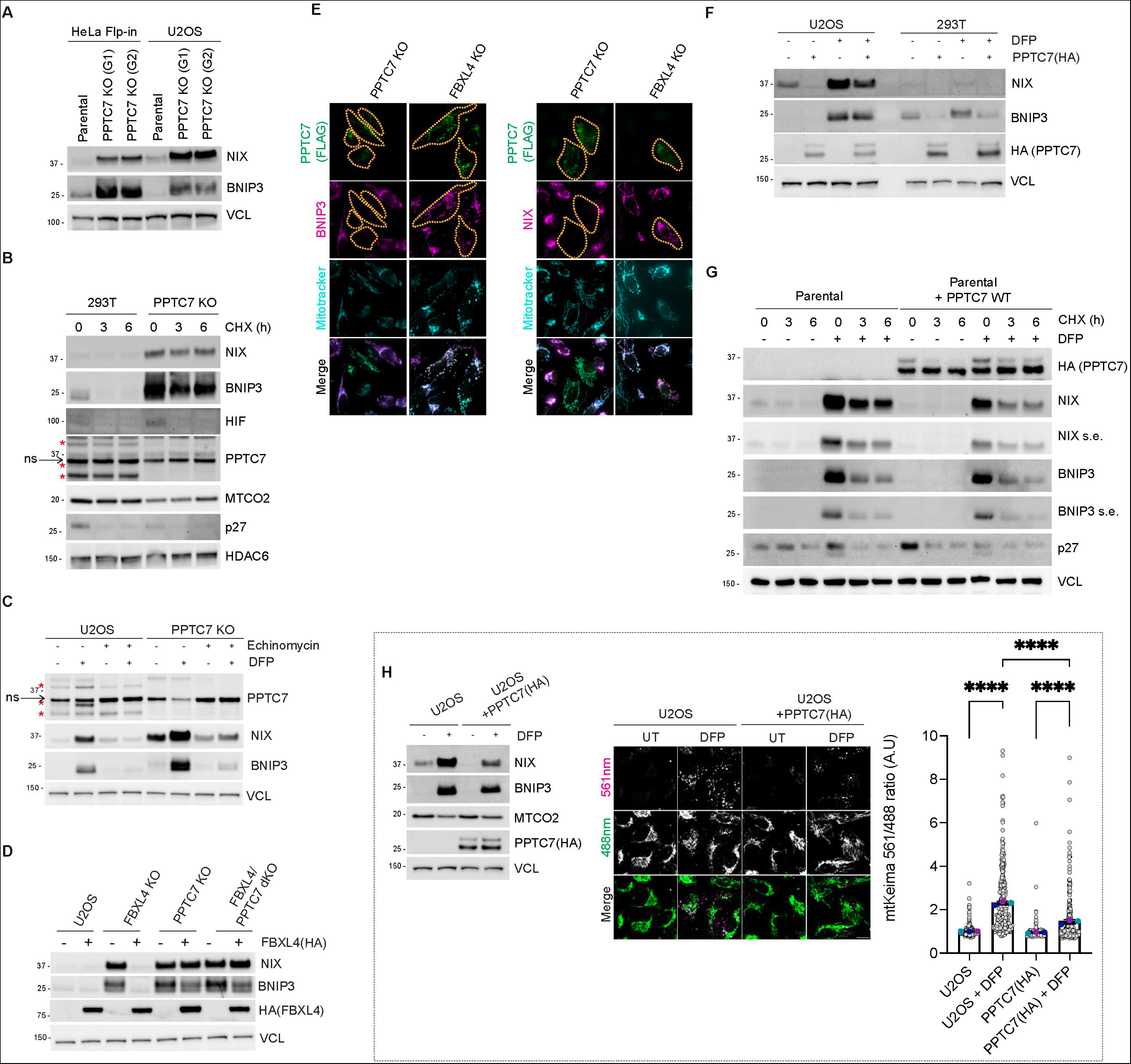
PPTC7 is required for the FBXL4-mediated destabilization of NIX/BNIP3. A. *Distinct guide RNAs targeting PPTC7 result in BNIP3 and NIX upregulation*. Cells were transfected PPTC7 guide 1 (G1) or guide 2 (G2). The red asterisks indicate PPTC7 specific bands and 28 kDa, 32 kDa, and 40 kDa. ns=non-specific band at ∼36 kDa. B. *BNIP3 and NIX are stabilized in PPTC7-deficient 293T cells.* Cells were treated with cycloheximide for the indicated times before immunoblotting as indicated. C. *Inhibition of HIF1*α *with echinomycin does not prevent the accumulation of BNIP3 and NIX in PPTC7-deficient cells.* U2OS cells or PPTC7-KO cells were treated with echinomycin for 24 hours. Echinomycin completely prevented the DFP-induced upregulation of BNIP3 and NIX, but only partially prevented the accumulation of BNIP3 and NIX in PPTC7 KO cells. DFP induces the 32 kDa form of PPTC7. D. *FBXL4 requires PPTC7 for its ability to promote BNIP3 and NIX turnover.* FBXL4-HA was expressed in parental, FBXL4 KO, PPTC7 KO, and FBXL4/PPTC7 dKO cells. BNIP3 and NIX protein levels were monitored by Western blotting in response to FBXL4 expression. E. *PPTC7-mediated downregulation of BNIP3 and NIX does not occur in FBXL4-deficient cells.* PPTC7-FLAG was transfected into either PPTC7 KO or FBXL4 KO cells. Cells were fixed and stained for FLAG(PPTC7) (green) and either NIX or BNIP3 (magenta). The orange dotted line surrounds the cells that have been transfected with PPTC7. F. *PPTC7(HA) overexpression causes the downregulation of BNIP3 and NIX in U2OS and 293T cells in basal conditions and after DFP treatment.* PPTC7(HA) was transduced into U2OS or 293T cells and the levels of BNIP3 and NIX were monitored by immunoblotting. G. *PPTC7 overexpression results in the destabilization of BNIP3 and NIX in basal conditions as well as after DFP treatment.* U2OS cells or U2OS cells stably transfected with PPTC7(HA) were treated with DFP for 24 hours. Cells were subjected to cycloheximide chase. s.e. = shorter exposure. H. *PPTC7 overexpression suppresses DFP-induced mitophagy.* Mitophagy was assessed U2OS mtKeima cells or U2OS mtKeima cells overexpressing PPTC7(HA) in the presence or absence of DFP. Emission signals at neutral pH were obtained after excitation with the 458 nm laser (green), and emission signals at acidic pH were obtained after excitation with the 458 nm laser 561 nm laser (magenta). Mitophagy is represented as the ratio of mt-Keima 561 nm fluorescence intensity divided by mt-Keima 458 nm fluorescence intensity for individual cells normalised to the mean of the untreated U2OS cells. Translucent grey dots represent measurements from individual cells. Coloured circles represent the mean ratio from independent experiments. The centre lines and bars represent the mean of the independent replicates +/- standard deviation. P values were calculated based on the mean values using a one-way ANOVA (*P<0.05, **P<0.005, ***P<0.001, ****P<0.0001). N=3.

**Supplementary Figure 2.**
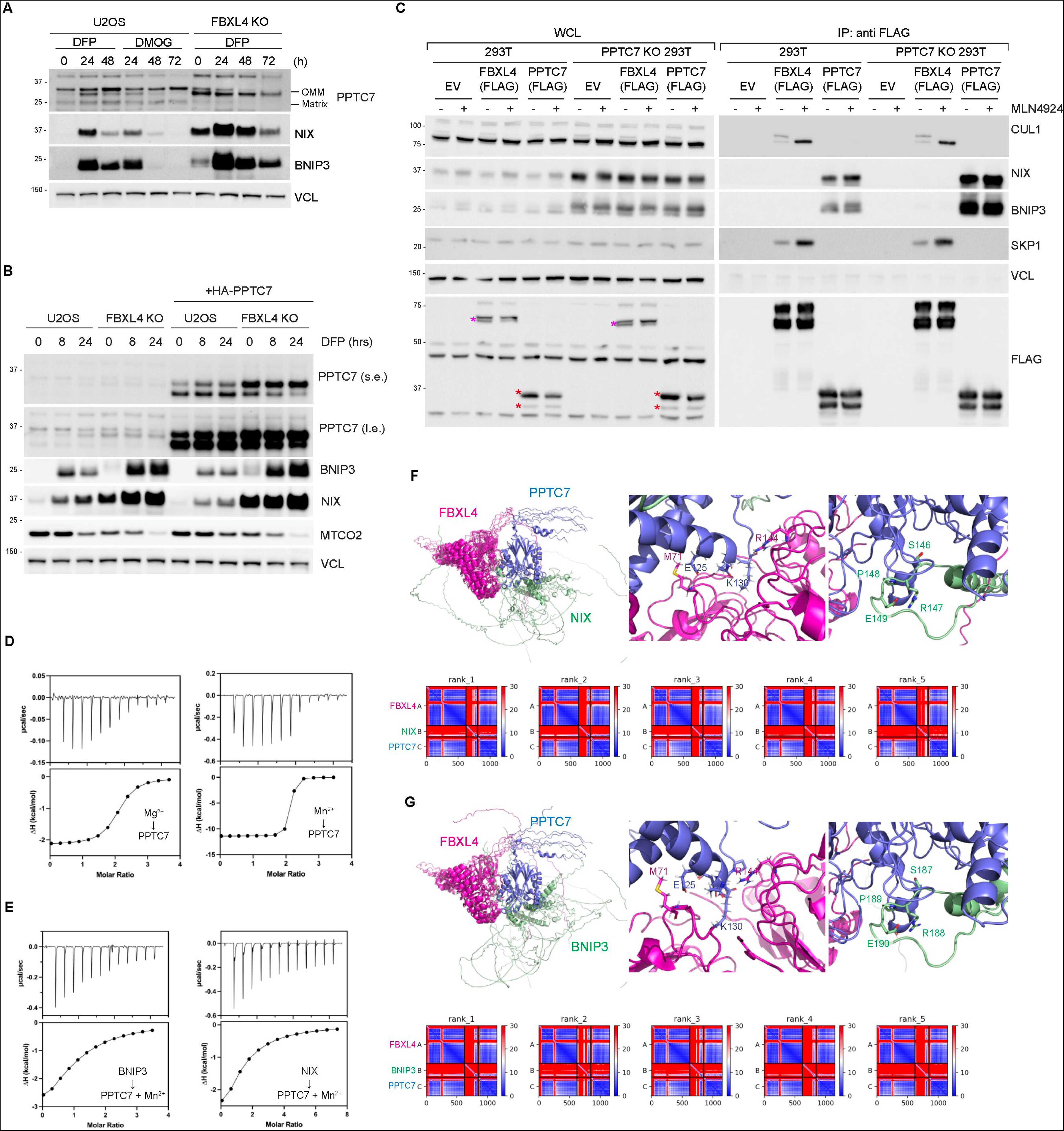
PPTC7 interacts with NIX/BNIP3 and FBXL4 and is not required for FBXL4 to interact with CUL1 and SKP1. A. Analysis of the stability of the outer membrane form of PPTC7 in response to DFP treatment, DMOG treatment, and/or FBXL4 deficiency. B. Analysis of the stability of the outer membrane form of endogenous PPTC7 and exogenous PPTC7(HA) in response to DFP treatment. C. *PPTC7 is not required for FBXL4 to interact with CUL1 or SKP1.* 293T or 293T PPTC7 KO cells were transfected with either FLAG-FBXL4 or FLAG-PPTC7. Cells were treated with MLN4924 for 24 hours where indicated. Cell lysates were immunoprecipitated with anti-FLAG beads, and the immuno-precipitates were analysed by immunoblotting as indicated. FBXL4 binds to CUL1 and SKP1, whereas PPTC7 binds to BNIP3 and NIX. WCL = whole cell lysates. D. ITC comparison of PPTC7 wild-type binding to Mg^2+^ and Mn^2+^. The binding affinities were 6.21 nM for Mn^2+^ and 237 nM for Mg^2+^. E. ITC comparison of PPTC7 wild-type binding to BNIP3 and NIX in the presence of Mn^2+^. The binding affinities were 20.1 μM for BNIP3 and 37.5 μM for NIX. F. Overlay of the top 5 AlphaFold2 models of FBXL4, PPTC7, NIX with predicted aligned error (PAE) plots. G. *Overlay of the top 5 AlphaFold2 models of FBXL4, PPTC7, BNIP3 with PAE plots*.

**Supplementary Figure 3.**
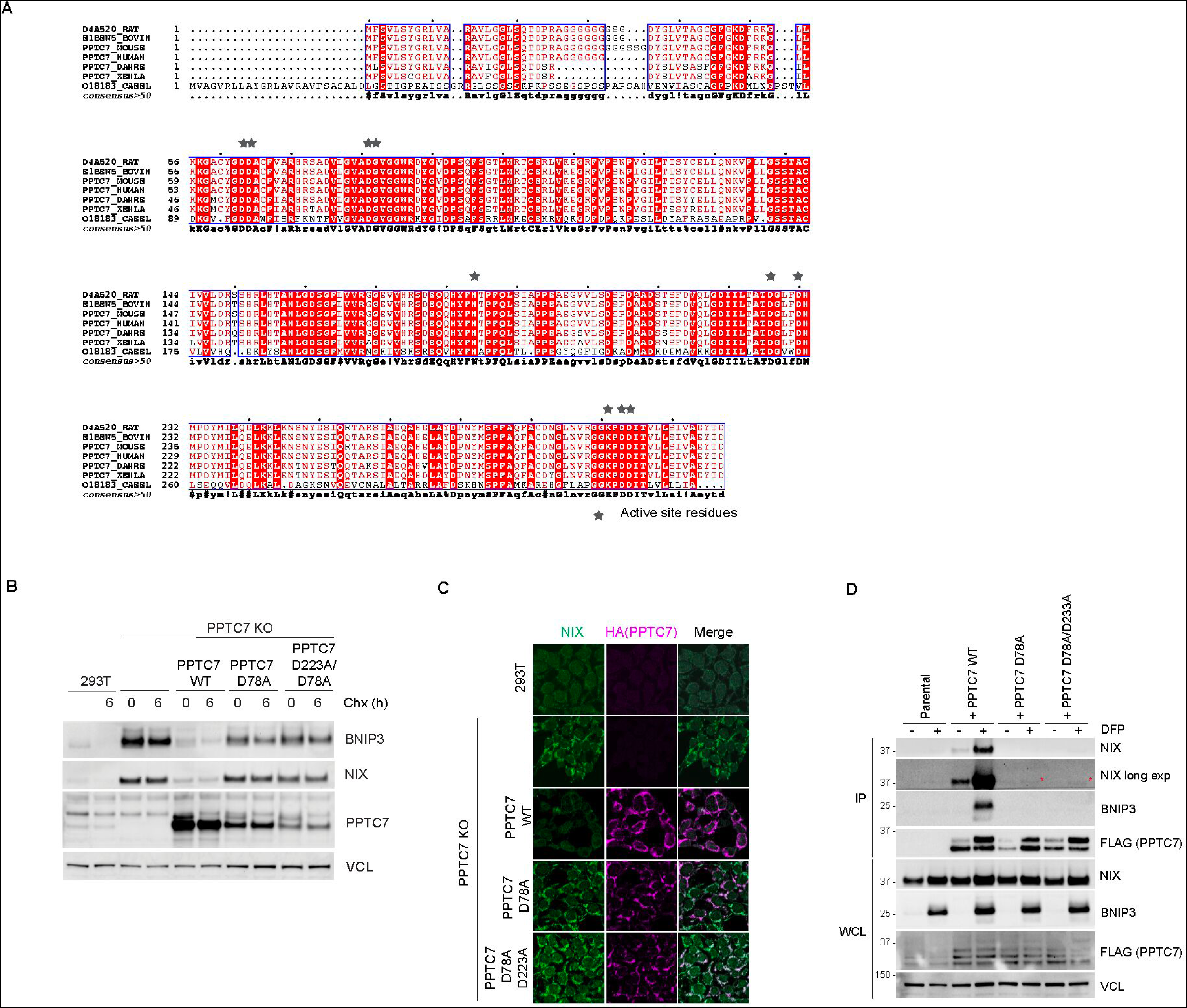
Radical disruption of PPTC7’s catalytic site interferes with its binding to BNIP3 and NIX. A. *Sequence alignment of PPTC7 orthologues with active site residues indicated*. B. *Disruption of the PPTC7’s active site residues from aspartate to alanine compromises its ability to downregulate BNIP3 and NIX.* PPTC7 KO cells were transduced with PPTC7 wild-type, PPTC7-D78A, or PPTC7-D223A/D78A. Wildtype PPTC7 rescued the turnover of BNIP3 and NIX, however, the D78A and D223A/D78A variants did not. C. *Aspartate to alanine mutations in PPTC7’s active cannot rescue the downregulation of NIX by PPTC7.* PPTC7 KO cells were complemented with PPTC7 or active site mutants and NIX levels were analysed by immunofluorescence microscopy. D. *Disruption of the PPTC7’s active site residues from aspartate to alanine interferes with its ability to bind to BNIP3 and NIX.* Cell lysates were immunoprecipitated with anti-FLAG beads, and the immuno-precipitates were analysed by immunoblotting as shown. Unlike the D78N mutant in Figure 3B, the D78A mutant is unable to bind to BNIP3 or NIX.

**Supplementary Figure 4.**
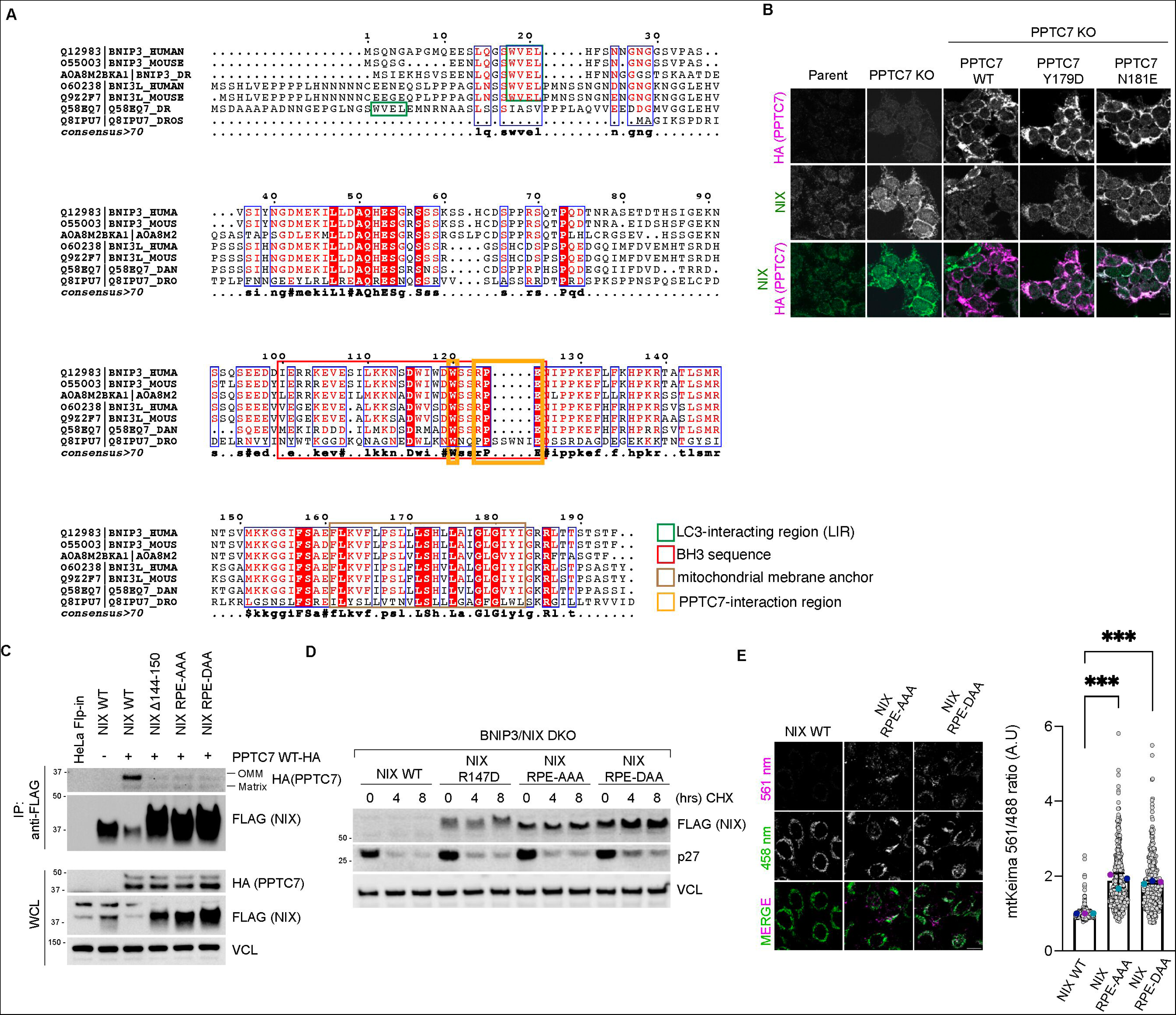
The NIX-PPTC7 interaction is critical for NIX turnover and mitophagy suppression. A. *Sequence alignment of BNIP3 and NIX orthologues.* Functionally relevant motifs or domains are indicated. B. *PPTC7-Y179D and PPTC7-N181E variants localise to mitochondria.* Wild-type PPTC7 can reduce NIX levels when expressed in PPTC7 KO cells, but PPTC7-Y179D and PPTC7-N181E cannot. C. *Arg147 in NIX is critical for binding to PPTC7.* PPTC7(HA) was transduced into cell lines expressing inducible NIX mutants. NIX expression was induced with doxycycline for 24 hours. Cell lysates were immunoprecipitated with anti-FLAG beads, and the immuno-precipitates were analysed by immunoblotting. D. *Arg147 in NIX is critical for its turnover.* HeLa Flp-in BNIP3/NIX double KO cells expressing FLAG-tagged NIX WT or NIX binding mutants (FLAG-tagged NIX Δ144-150, NIX RPE-AAA and NIX RPE-DAA) were subjected to a cycloheximide chase. E. *Expression of NIX-RPE/AAA and NIX-RPE/DAA in NIX leads to an increase in basal levels of mitophagy compared with NIX-wildtype.* Hela Flp-In NIX knockout/ Hela Flp-In BNIP3/NIX double knockout Keima cells stably expressing NIX mutants were treated with doxycycline for 48 hours and mitophagy was evaluated using live-cell confocal fluorescence microscopy.

## Methods and Materials

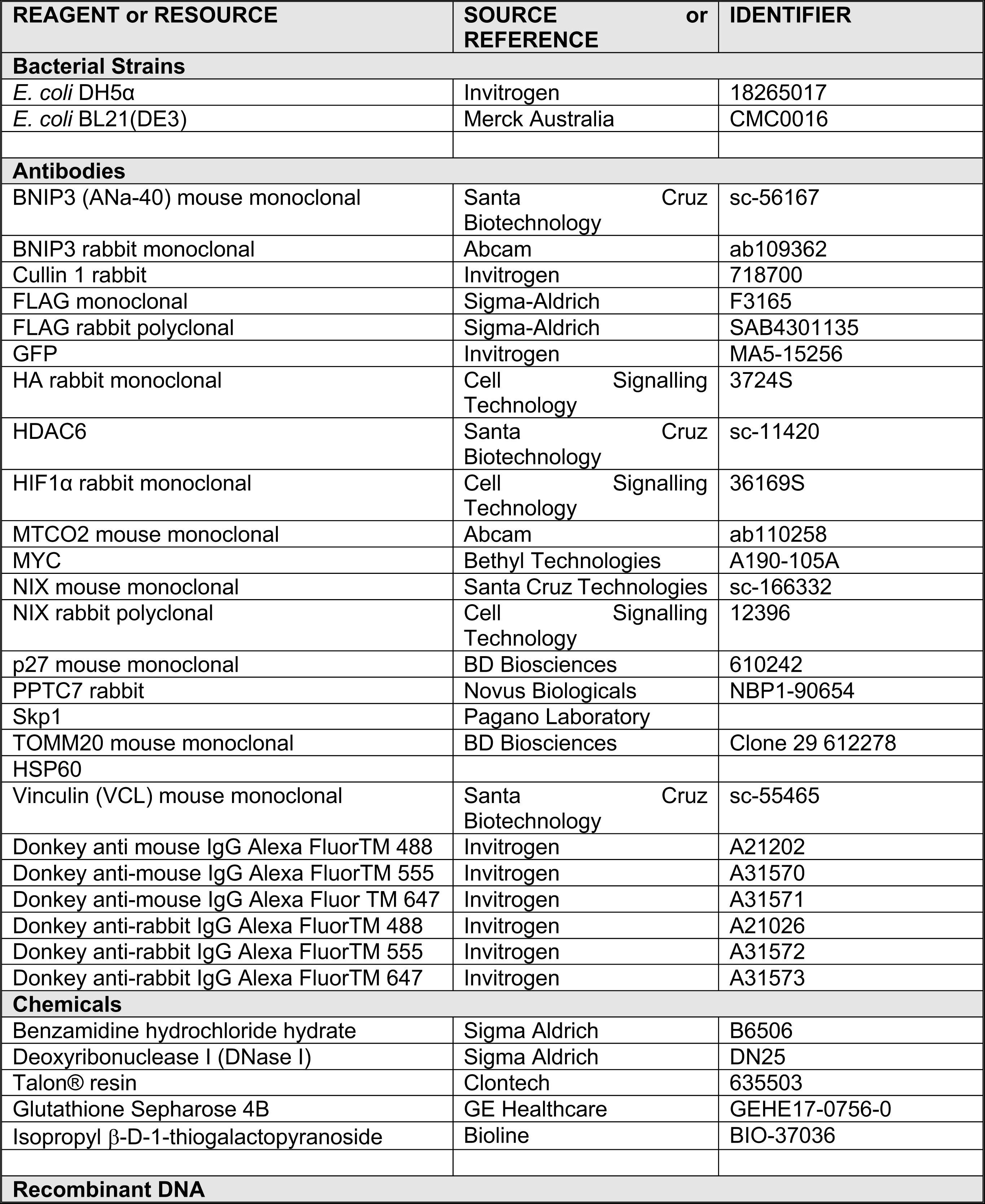
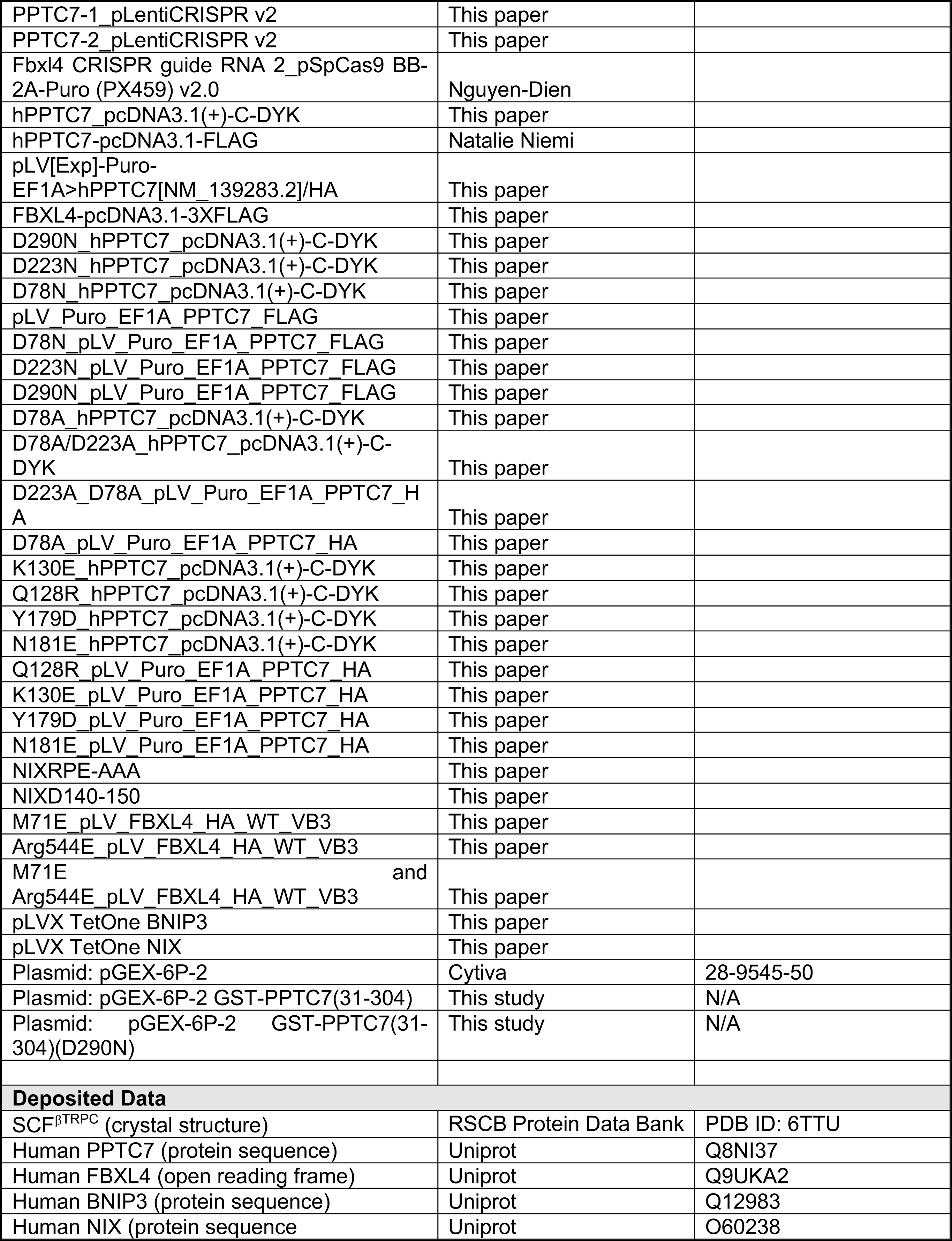
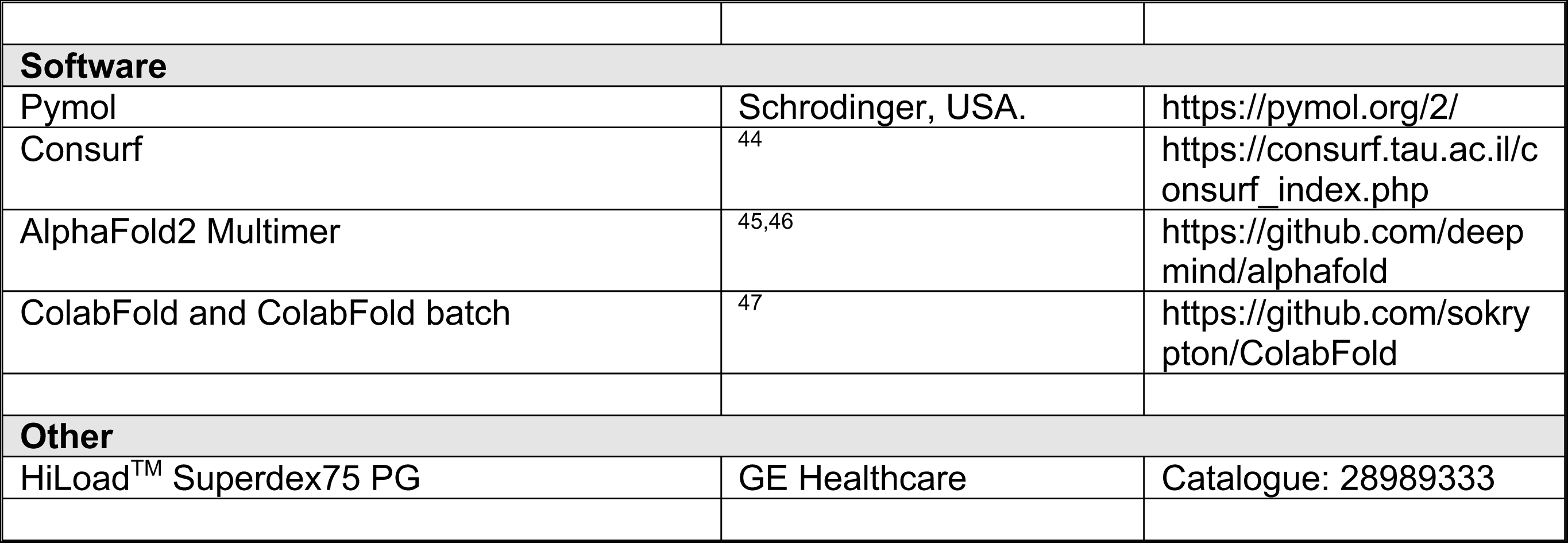

### CRISPR-CAS9 mediated genome editing

To generate HeLa and U2OS PPTC7 KO cell lines, PPTC7-1_pLentiCRISPR V2 and PPTC7-2_pLentiCRISPR V2 plasmids were generated by Genscript^®^ based on the the following gRNA sequences: TTCGTACCTAGTAATCCCAT and CGGCGACTACGGACTGGTGA, respectively. The pLenti-CRISPR V2 plasmids were used to generate lentiviruses in 293T cells. HeLa or U2OS cells were transduced with lentivirus, and approximately 24 hours post-transduction, cells were selected with puromycin for 48 hours. Successful knockout was confirmed using immunoblotting using antibodies to PPTC7. FBXL4 deficient U2OS cells (clone 2G10) have been described previously ^18^. PPTC7 deficient 293T cells have been described previously ^48^.

### Cell culture, transfection, and chemicals

Cell lines were incubated at 37°C in a humidified incubator containing 5% CO_2_. HeLa (ATCC CCL-2), U2OS (ATCC HTB-96) and HEK293T (ATCC CRL-3216) cells were propagated in Dulbecco’s Modified Eagle’s medium/Nutrient mixture F-12 GlutaMAX™ (DMEM/F-12; Thermo Fisher Scientific) supplemented with 10% foetal bovine serum (Gibco). All cell lines were regularly screened for mycoplasma contamination. Plasmid transfections were performed using Lipofectamine 2000 (Thermo Fisher Scientific) according to the manufacturer’s recommendations. Cells were transfected with plasmid DNA using Lipofectamine 2000 (Thermo Scientific) for approximately 48 h. The following chemicals from Sigma were used: cycloheximide (CHX; 100 μg/ml; 66-81-9), deferiprone (DFP; 1 mM; 379409), DMOG (0.5 mM; D3695) and echinomycin (10 nM; SML0477). MLN4924 (0.5 μM; 85923S) was obtained from Cell Signaling Technology. MG132 (10 μM; 474787) was purchased from Merck.

HeLa mtKeima cell lines expressing dox-inducible FLAG-S tag NIX-wild-type and NIX mutants were generated as described previously ^49^. Briefly, pcDNA5/FRT/TO (Thermo Fisher) constructs expressing NIX or variants were co-transfected with pOG44 into HeLa-T-rex Flp-in cells mtKeima cells to generate inducible cell lines using Flippase (Flp) recombination target (FRT)/Flp-mediated recombination technology. Twenty-four hours post-transfection, cells were selected with Hygromycin B (400 μg/ml) for approximately 10 days. To induce expression, cells were treated with 0.5 μg/ml doxycycline (Sigma; 10592-13-9).

### Virus production and transduction

Lentiviruses (pLV constructs) or retroviruses (pCHAC-mt-mKeima) were packaged in HEK293T cells. The media containing lentiviral or retroviral particles were harvested 48 h later. Cells were transduced with virus along with 10 μg/mL polybrene (Sigma). Following transduction, cell lines were either sorted using FACS based on fluorescence (for mtKeima) or selected with puromycin (for cells with pLV constructs).

### Biochemical techniques, immunoblotting, and co-immunoprecipitation

Immunoblotting was performed as previously described ^18^. Cultured cells were harvested and lysed in SDS lysis buffer (50 mM Tris, 2% SDS) followed by heating at 95°C for 15 minutes. Protein extracts were diluted in Bolt™ LDS Sample Buffer (Invitrogen™; B0008). Equal amounts of protein samples were separated using BOLT pre-cast 4–12% gradient gels (Invitrogen™) and transferred onto methanol-pretreated Immobilon^®^-P PVDF Membrane (0.45 μm pore size) (Merck; IPVH00010) using BOLT gel transfer cassettes and BOLT transfer buffer (Invitrogen™; BT0006). Membranes were blocked in 5% skim milk for 1 h at room temperature followed by overnight incubation at 4°C with the indicated primary antibodies. Chemilumiscent detection of HRP-conjugated secondary antibodies was performed using Pierce ECL Western blotting substrate (Thermo Fisher Scientific; 32106) or Pierce SuperSignal West Femto Substrate (Thermo Fisher Scientific; 34094) and ChemiDoc™ Imaging System (Bio-Rad). Phos-tag Precast gels were purchased from Fujifilm/WAKO and run according to the manufacturer’s instructions. For immunoprecipitation, cellular lysis was performed using a Tris-Triton lysis buffer (50 mM Tris-Cl pH 7.5, 150 mM NaCl, 10% glycerol, 1 mM EDTA, 1 mM EGTA, 5 mM MgCl2, 1 mM β-glycerophosphate, and 1% Triton), supplemented with protease inhibitor cocktail (Rowe Scientific; CP2778) and PhosSTOP EASYpack Phosphatase Inhibitor Cocktail (Roche; 4906837001), and kept on ice for 30 minutes. Subsequently, cell lysates were centrifuges at 21,130 g for 10 minutes at 4°C. For immunoprecipitation of exogenously expressed FLAG-tagged or HA-tagged proteins, the cell lysates were incubated with bead-conjugated FLAG (Sigma; A2220) or bead-conjugated HA (Thermo Fisher Scientific; 88837) respectively, in a rotating incubator for 1-2 hours at 4°C. The immunoprecipitated complexes were then washed five times with Tris-Triton lysis buffer before elution with Bolt™ LDS Sample Buffer for subsequent immunoblotting.

### Protease import protection assay

Crude mitochondria were prepared by resuspending cell pellets in mitochondrial isolation buffer (250 mM mannitol, 0.5 mM EGTA and 5 mM HEPES–KOH pH 7.4) followed by homogenisation using a 26.5 G needle (303800, Becton Dickinson) for 10 strokes, as in ^50^. The homogenate was then centrifuged twice at 600 g for 10 minutes at 4 °C to remove cell debris and intact nuclei. The supernatant was then centrifuged twice at 7000 g for 10 minutes at 4 °C to acquire a mitochondrial pellet. The resuspended pellets were centrifuged twice at 10 000 g for 10 minutes at 4 °C. Isolated mitochondria were then divided into untreated or proteinase K treated conditions (10 μg/ml Proteinase K, EO0492, Life Technologies). Samples were rotated for 20 minutes at 4 °C. Proteinase K digestion was then blocked with 1 mM phenylmethylsulfonyl fluoride (PMSF, sc-482875, Santa Cruz Biotechnology) on ice for 10 min. Samples were centrifuged at 10 000 g for 10 minutes at 4 °C. Pellets were resuspended in 1X NuPAGE LDS sample buffer (NP0007, Life Technologies) and analysed by immunoblotting.

### Protein expression and purification

GST-PPTC7 (31-304) in the pGEX-6P-2 vector was transformed into BL21(DE3) cells and plated on LB-Agar plates containing 0.1 mg/mL ampicillin. A single colony from this plate was used to inoculate 200 mL of LB medium (containing 0.1 mg/mL ampicillin) and grown overnight at 37°C with shaking at 180 rpm. The following day, 10 mL of overnight culture was used to inoculate 1L of LB medium, supplemented with 0.1 mg/mL ampicillin. Cultures were grown at 37°C with shaking at 190 rpm until the OD_600_ reached 0.9-1.0, at which point protein expression was induced by adding 0.5 mM IPTG and the temperature was lowered to 20°C. Cultures continued to grow overnight. Cells were harvested the following day by centrifugation at 6,200 rpm for 7 minutes using a Beckman JLA 8.1000 rotor. Cells were resuspended in lysis buffer (50 mM Tris, pH 8.0, 500 mM NaCl, 10% glycerol, benzamidine (0.1 mg/mL) and DNase (0.1 mg/mL) and lysed using a Cell Disruptor (Constant Systems) at 35 kPsi. The cell lysate was centrifuged at 14,000 rpm for 40 minutes at 4°C using a Beckman JLA 16.250 rotor. The soluble fraction was then purified by affinity chromatography using glutathione-sepharose resin. The GST tag was removed from the protein by incubating the resin with PreScission protease overnight at 4°C. Untagged PPTC7 was eluted the following day in lysis buffer. The elution was further purified by gel filtration chromatography using a Superdex 75 (16/600) column (GE Healthcare) into buffer containing 50 mM HEPES (pH 7.5), 300 mM NaCl and 2 mM β-mercaptoethanol.

### Isothermal titration calorimetry (ITC)

All ITC experiments were performed using a Microcal PEAQ-ITC instrument (Malvern) at 25°C. Before performing the metal-binding experiments, PPTC7 was incubated with 1 mM EDTA and then gel-filtered using a PD10 desalting column (Cytiva) to remove any bound metal ions. All metal-binding experiments were performed in buffer containing 50 mM HEPES (pH 7.5), 200 mM NaCl and 2 mM β-mercaptoethanol. In these experiments, 150 µM MnCl_2_ or MgCl_2_ was titrated into 15 µM PPTC7. Experiments to test the binding of PPTC7 to BNIP3 and NIX peptides were performed in buffer containing 50 mM HEPES (pH 7.5), 300 mM NaCl, 10 mM MnCl_2_ and 2 mM β-mercaptoethanol. In all experiments, 500 µM of peptide was titrated into 25 µM PPTC7. The dissociation constant (K_D_), enthalpy of binding (ΔH) and stoichiometry (N) were calculated using the MicroCal PEAQ-ITC software.

### pNPP phosphatase assay

The generic phosphatase substrate para-nitrophenyl phosphate (pNPP) was used to determine the phosphatase activity of recombinant PPTC7 wild-type and PPTC7-D290N mutant. 10 mM pNPP (New England Biolabs) was added to 4 µM PPTC7 in the presence or absence of either 5 mM MnCl_2_ or 5 mM MgCl_2_. All reactions were carried out in buffer containing 50 mM HEPES (pH 7.5), 200 mM NaCl and 2 mM β-mercaptoethanol, in a total volume of 100 µL. The reaction was incubated for 15 minutes at room temperature before measuring the absorbance at 405 nm using an Infinite M1000 Pro plate reader (Tecan).

### Protein structural prediction, modelling and visualisation

All protein models were generated using AlphaFold2 Multimer ^45,46^ implemented in the Colabfold interface available on the Google Colab platform^47^. For each modelling experiment ColabFold was executed using default settings where multiple sequence alignments were generated with MMseqs2^51^. For all final models displayed in this manuscript, structural relaxation of peptide geometry was performed with AMBER^52^. For all modelling experiments, we assessed (i) the prediction confidence measures (pLDDT and interfacial iPTM scores), (ii) the plots of the predicted alignment errors (PAE) and (iii) backbone alignments of the final structures. The model of PPTC7 and NIX bound to the SCF^FBXL4^ complex was constructed in two steps. Firstly, the trimeric PPTC7–NIX–FBXL4 complex was predicted with AlphaFold2 as described above. This was then aligned to the structure of SCF^βTRCP^ consisting of the proteins βTRCP–Skp1– Cul1–Rbx1–Ube2d–Nedd8–Ub (PDB ID 6TTU)^53^. All structural images were made with Pymol (Schrodinger, USA; https://pymol.org/2/).

### Indirect immmunofluorescence staining and mtKeima assay

Cells grown as monolayers on coverslips were fixed with ice-cold methanol. Cells were blocked with 2% BSA in PBS. Cells were then sequentially labelled with primary antibodies for 1 h, followed by the species-specific secondary antibodies for 1 h. Coverslips were mounted on glass microscope slides using Prolong Diamond Antifade Mountand (Thermo Fisher Scientific; P36965). Images were acquired using either a DeltaVision Elite inverted microscope system (GE Healthcare) or a using a ×60/1.4NA Oil PSF Objective from Olympus or Zeiss LSM900 Fast AiryScan2 Confocal microscope with a 63× C-Plan Apo NA 1.4 oil-immersion objective. DeltaVision images were processed using the Softworx deconvolution algorithm whereas Airyscan images were processed using ZEN Blue 3D software (version 3.4).

The mt-Keima assay was performed as previously described (Sun et al., 2017). A Leica DMi8 SP8 Inverted confocal microscope equipped with a 63x Plan Apochromatic objective and environmental chamber (set to 5% CO2 and 37°C) was used to capture images. Quantitative analysis of mitophagy with mt-Keima was performed using Image J/Fiji software. Individual cells were isolated from the field of view by creating regions of interest (ROI). The chosen ROI were then cropped and separated into distinct channels before undergoing threshold processing. The fluorescence intensity of mt-Keima at 561 nm (indicating lysosomal signal) and 458 nm (indicating mitochondrial signal) at the single-cell level was measured, and the ratio of 561 nm to 458 nm was calculated. Three biological replicates were performed for each experiment, with >50 cells analyzed per condition for each repeat.

### Statistical Analysis

GraphPad Prism 9.0 software was used to perform statistical comparisons. The centre line and error bars on the graphs represent the mean and standard deviation of normalised biologically independent replications. Unless otherwise noted, three or more biologically independent replications for used for statistical comparisons. No blinding or randomisation was incorporated into the experimental design. *P* values greater than 0.05 were considered non-significant.

## Acknowledgements

We thank Natalie Niemi and David Pagliarini for helpful discussions, the FLAG-PPTC7 plasmid, and the PPTC7 KO 293T cells. Imaging was performed at the Microscopy and Image Analysis Facility in the School of Biomedical Sciences at the University of Queensland. Lentiviruses were produced by the University of Queensland (UQ)-Viral Vector Core. This work was supported by the Australian National Health and Medical Research Council (APP1183915 and APP1136021), a Brain Foundation Research grant (2020), a Mito Foundation Incubator Grant (2022), Mito Foundation Research Fellowship to P.K. and a Mito Foundation Scholarship Top-up to K.L.K. and an Australian Research Council Future Fellowship (FT180100172) to J.K.P. B.M.C is supported by an Australian NHMRC Investigator Grant (APP2016410).

